# Entropy Sorting Feature Selection: information-theoretic gene set identification improves single-cell RNA sequencing data interpretability

**DOI:** 10.64898/2026.01.26.701684

**Authors:** Arthur Radley, Giulia Boezio, Cameron Shand, Ruben Perez-Carrasco, James Briscoe

**Affiliations:** The Francis Crick Institute, Developmental Dynamics Laboratory, London, UK; The Francis Crick Institute, Software Engineering & AI STP, London, UK; Imperial College London, Department of Life Sciences, London, UK

## Abstract

Single-cell RNA sequencing (scRNA-seq) has transformed our ability to resolve cellular heterogeneity, but extracting meaningful signals remains challenging due to technical noise and batch effects. Most methods for denoising scRNA-seq data have focused on using latent representations such as principal component analysis and deep learning to prioritise biological signals. By contrast, despite its influence on downstream analyses, feature selection has received relatively limited attention, leading to widespread reliance on the comparatively simplistic strategy of highly variable gene selection. Here we present Entropy Sorting Feature Selection (ESFS), a modular, user-friendly framework that substantially improves the interpretability of scRNA-seq data. Notably, ESFS reveals complex expression dynamics that are obscured in latent representations. We demonstrate the utility of ESFS in diverse data: identifying coherent developmental programs across eight independent human embryo datasets without batch integration; resolving spatial gene expression in mouse colon missed by conventional analyses; disambiguating shared and tumour-specific microenvironments in glioblastoma; and disentangling spatial, temporal, and neurogenic programs in the developing mouse neural tube. Beyond delivering a powerful and user-friendly software that deepens insight into complex biological systems, our work establishes Entropy Sorting as a novel information theoretic for advanced data analysis methods.

## 1 Introduction

Single-cell RNA sequencing (scRNA-seq) has become a pervasive technique for investigating biological systems at cellular resolution. Although the number and variety of experimental scRNA-seq protocols have grown rapidly^1^, the core analytical objective remains unchanged: to identify informative gene expression patterns that reveal cellular identity and function. However, the high dimensionality of scRNA-seq data, coupled with technical noise and uncertain biological ground truths, continues to challenge the extraction of meaningful biological signals^2^.

Standard analysis pipelines such as Seurat and Scanpy address these issues by combining highly variable gene (HVG) selection with feature extraction/latent representation techniques such as principal component analysis (PCA). These workflows have become widely adopted because of their robustness and ease of use. Yet, they come with limitations: HVG selection often lacks reproducibility across different implementations^3^ and can introduce distortions and biases into data analysis^4,5^. Likewise, PCA or other dimensionality reduction methods can distort true biological variation by introducing artefactual structure^6^. Moreover, identification of cell type marker genes using conventional scRNA-seq analysis workflows is typically tied to enrichment of expression in discrete clusters of cells^7–9^ or genes^10^. As a result, these approaches can inadvertently mask biologically meaningful gene expression heterogeneity or complex hierarchical patterns that span combinations of discrete clusters. To fully realise the potential of scRNA-seq data, new methods are needed that maximise data interpretability while minimising the risk of introducing computational artefacts.

Because information-theoretic measures such as mutual information can non-parametrically quantify dependencies between genes without assuming a specific underlying statistical model, they provide an attractive framework for identifying patterns of gene co-expression. However, estimating mutual information directly from non-discretised scRNA-seq expression profiles is often statistically unreliable and can become prohibitively computationally expensive in high-dimensional settings^11,12^. In addition, unlike classical correlation measures, mutual information does not admit a simple closed-form null distribution; statistical significance is therefore typically assessed using permutation or bootstrap procedures that further add to the computational burden when applied at scale across large numbers of gene pairs^13,14^.

In previous work, we introduced Entropy Sorting (ES), a novel information-theoretic framework that addresses both the accuracy and computational limitations of conventional information-theoretic approaches^5^. Application of ES to scRNA-seq data has enabled the identification of fine-grained developmental trajectories across diverse datasets^5,15–21^. However, theoretical gaps in the original framework limited its generalisability. Here, we rigorously formalise and extend the ES mathematical foundation to robustly quantify gene–gene dependencies. Building on these advances, we introduce a modular software suite, Entropy Sorting Feature Selection (ESFS), which integrates three ES-based algorithms (Fig 1) to address persistent challenges in scRNA-seq analysis.

**Figure 1:**
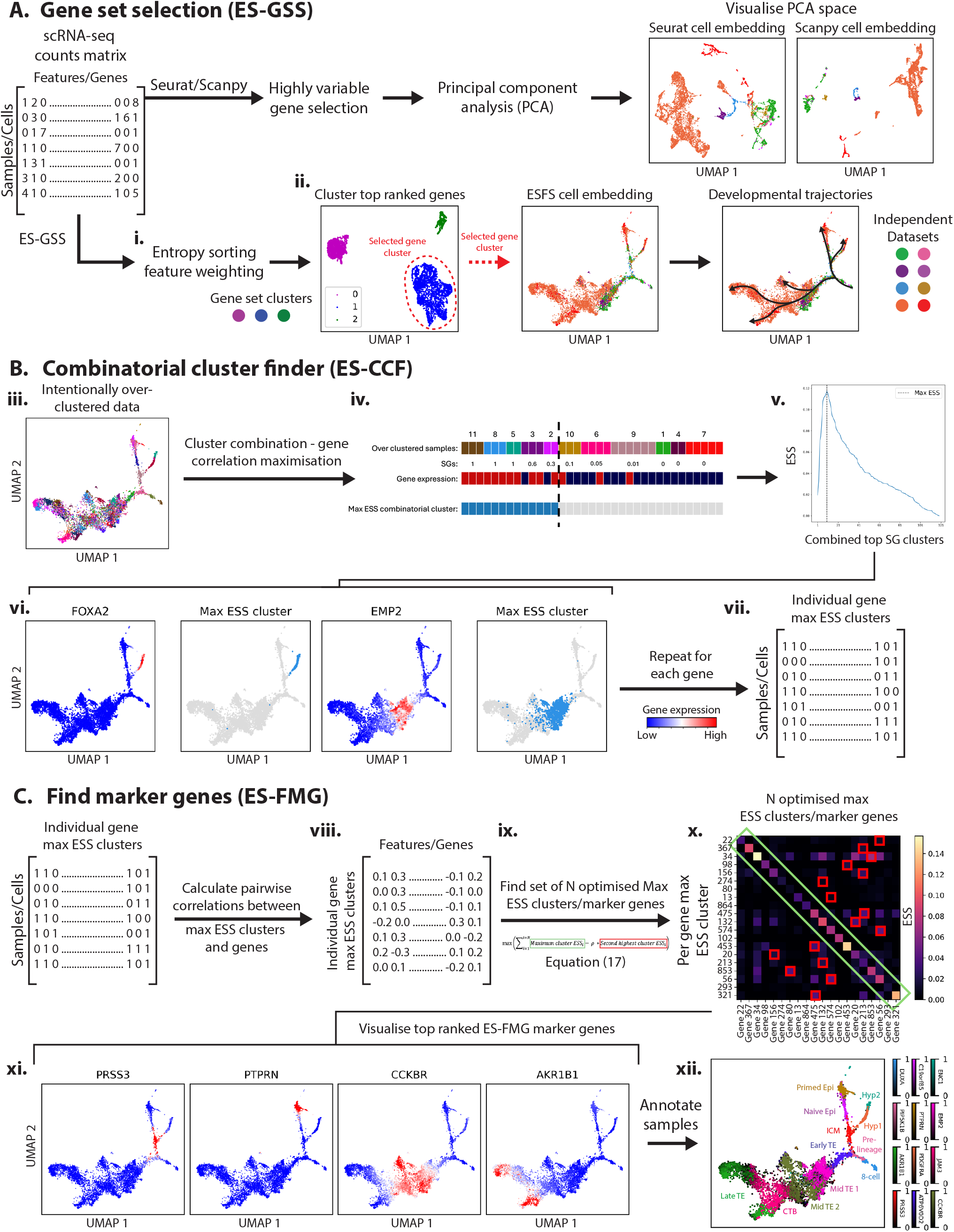
Core ESFS algorithms: The ESFS workflow comprises three packages. **A**. ES-Gene Set Selection (ES-GSS) separates technical noise from biological signals of interest. The approach reveals complex gene expression dynamics without using computational batch integration software that may unintentionally introduce computational artefacts (see Fig 2 for more detail). **B**. To remove the ambiguity regarding which resolution of sample clustering best identifies gene expression profiles of interest in a given dataset, ES-Combinatorial Cluster Finder (ES-CCF) takes intentionally over-clustered data and identifies combinations of clusters that maximise correlation with individual genes. **C**. To simplify robust marker gene identification, ES-Find Marker Genes (ES-FMG) uses the output of ES-CCF to solve an optimisation problem to identify a set of *N* marker genes that capture distinct and robust gene expression profiles present in the data. For further details of i-xii see Materials and Methods.

The first module, ES-Gene Set Selection (ES-GSS) (Fig 1A), provides a principled alternative to highly variable gene (HVG) selection by enabling robust multivariate feature selection. Rather than ranking genes based solely on univariate dispersion, ES-GSS applies Entropy Sorting to non-parametrically identify genesets exhibiting significant co-expression structure. Across multiple datasets, we show that ES-GSS improves interpretability by resolving cellular states and developmental trajectories that are often conflated under standard HVG-based workflows and latent-space representations. Moreover, when applied to datasets generated by independent studies, ES-GSS-based feature selection outperforms conventional integration approaches, enhancing data interpretability while reducing the risk of introducing computational artefacts^22^.

A second challenge in scRNA-seq analysis is choosing an appropriate clustering resolution that accurately captures both broad and fine-grained expression patterns^23^. Most existing pipelines require users to fix a discrete number of clusters prior to downstream steps such as differential gene expression analysis. This creates a trade-off: too few clusters can mask important sub-populations, while too many risk over-partitioning cells into artificial groups that lack molecular distinction.

To overcome this, we developed ES-Combinatorial Cluster Finder (ES-CCF) (Fig 1B). ES-CCF transforms a previously intractable combinatorial clustering problem—one that would require longer than the age of the universe to exhaustively evaluate—into a linear-time procedure that can be solved in minutes on a standard laptop. ES-CCF identifies sets of cells in which specific genes are maximally co-expressed. This reveals both fine-scale and broad-scale gene expression profiles without being constrained by a fixed clustering resolution.

The third challenge stems from a fundamental mismatch: gene expression varies continuously across cells, but most analytical tools impose discrete, non-overlapping cluster boundaries. This disconnect limits our ability to identify minimal sets of marker genes that accurately capture robust expression patterns in both independent and overlapping cell states.

To address this, we developed ES-Find Marker Genes (ES-FMG) (Fig 1C), which identifies a compact set of marker genes that best distinguish expression programs across the combinatorial cluster space uncovered by ES-CCF. Unlike traditional marker selection approaches, combining ES-CCF with ES-FMG relaxes the constraints of relying on fixed sample clustering or assumptions of exclusivity, making it ideally suited for unsupervised, high-resolution marker discovery. Further, ES-FMG’s objective function seeks to identify a set of genes that show robust expression within distinct minimally overlapping cell populations, reducing redundancy in the optimised marker gene set. This is crucial for downstream applications such as combinatorial genetic perturbation or spatial targeting^24^.

Applied to datasets with distinct technical and biological challenges, ESFS consistently uncovers deeper and more structured biological insights than current state-of-the-art methods. In pre-implantation human embryo scRNA-seq data, ESFS reveals coherent developmental trajectories across datasets, without batch integration that can distort biological signals^6^. In mouse colon spatial transcriptomic data, ESFS identifies cell populations and gene expression dynamics missed by the widely used consensus Non-negative Matrix Factorisation (cNMF)^25^. Applied to glioblastoma spatial transcriptomic data, ESFS reveals global, shared, and tumour-specific expression programs within a single, easy-to-use workflow. Finally, applied to the developing mouse neural tube^26^, ESFS produces robust developmental trajectories and resolves distinct spatial, temporal, and differentiation gene expression programs that remain unresolved using conventional clustering-based methods. We experimentally validate that top-ranked marker genes identified by ESFS outperform those selected by differential expression in Seurat or Scanpy.

Our results demonstrate that ESFS offers a noise-resilient alternative to conventional scRNA-seq workflows, enabling deeper insights into the structure of complex cellular systems. By providing a user-friendly package with interpretable outputs, ESFS allows the identification of biologically meaningful signatures that are often overlooked by standard approaches.

## 2 Results

### Entropy Sorting to Capture Complex Gene Relationships

The Entropy Sorting (ES) framework is derived from the well known quantity of conditional entropy (CE), with the aim of providing non-parametric, noise and data sparsity resistant, model free quantification of gene-gene correlations and their significance^5^. There are four mathematically valid rearrangements of the CE equation (Entropy Sort Equation 1-4, ESE1-4; Fig S1B). Using only one ESE is insufficient for robust quantification of correlation and significance (Materials and Methods). We incorporate all four ESE rearrangements (ESE1-4; Fig S1B), following robust logic outlined in the Materials and Methods to select the appropriate formulation dynamically for any given feature pair (Fig S1C). Together, these provide a mathematically rigorous foundation for software implementation and downstream analysis.

We tested this framework on a range of transcriptomic datasets and compared it with current practices in the field.

### ESFS characterises early human embryo development across independent datasets

A central goal of single-cell RNA-sequencing (scRNA-seq) analysis is the identification of distinct cell types and gene expression programs. This task is complicated by the high dimensionality, sparsity, and batch variability that characterise most scRNA-seq datasets. These issues are particularly acute in rare or difficult-to-obtain samples – such as early human embryos – where limited cell numbers and cross-study heterogeneity compound analytical challenges.

For example, to date, fewer than 10,000 cells have been profiled from human embryos spanning embryonic days 3 to 14 (E3–E14), with data collected across multiple research groups^16,27–33^. This scarcity, along with inter-batch variation, makes it difficult to reconstruct developmental trajectories without introducing artefacts.

To address these challenges, recent efforts have focused on integrating datasets through latent variable models and batch correction algorithms^28,34–36^. While such methods improve alignment across batches, they risk obscuring biologically meaningful variation or introducing artefactual structure^37,38^.

In this context, we demonstrate that the Entropy Sorting Feature Selection (ESFS) framework can provide a powerful alternative to batch integration by amplifying informative gene expression without requiring feature extraction or latent space approximation. Rather than harmonising datasets through dimensionality reduction, ESFS identifies a subset of genes where biological variance outweighs technical noise, enabling joint analysis of multiple embryonic datasets directly in the gene expression space. Applying ESFS to eight independent human embryo datasets (E3–E14), we generate a unified high-resolution embedding that captures biologically relevant cell states and trajectories.

Because ESFS operates in the gene expression space, it offers interpretable results and simplifies downstream tasks such as query dataset projection without the need for complex model retraining or cross-study correction pipelines^34,35^. By overcoming these key limitations of current integration frameworks, ESFS enables more robust, artefact-free characterisation of early developmental processes.

#### ES-GSS enables artefact free high quality dataset integration

To assemble the most comprehensive scRNA-seq dataset of human embryos up to E14 to date, we compiled data from eight independent studies^16,27–33^. Using the ES-GSS algorithm from our ESFS workflow, we started by ranking gene importance, followed by clustering of highly ranked genes to identify sets of genes with significant co-expression profiles (Fig 2A, left panel). From these distinct genesets, we found a set of 2435 genes (Fig 2A, gene cluster 2) that produced a high-resolution UMAP embedding capturing early human embryo development at single-cell resolution (Fig 2A, right panel). Notably, this embedding was generated directly from the gene expression space (i.e., without applying PCA for dimensionality reduction). The resulting UMAP shows clear dataset intermixing (Fig 2A), with evident cell-state bifurcations and logical temporal progression across developmental stages (Fig 2B).

**Figure 2:**
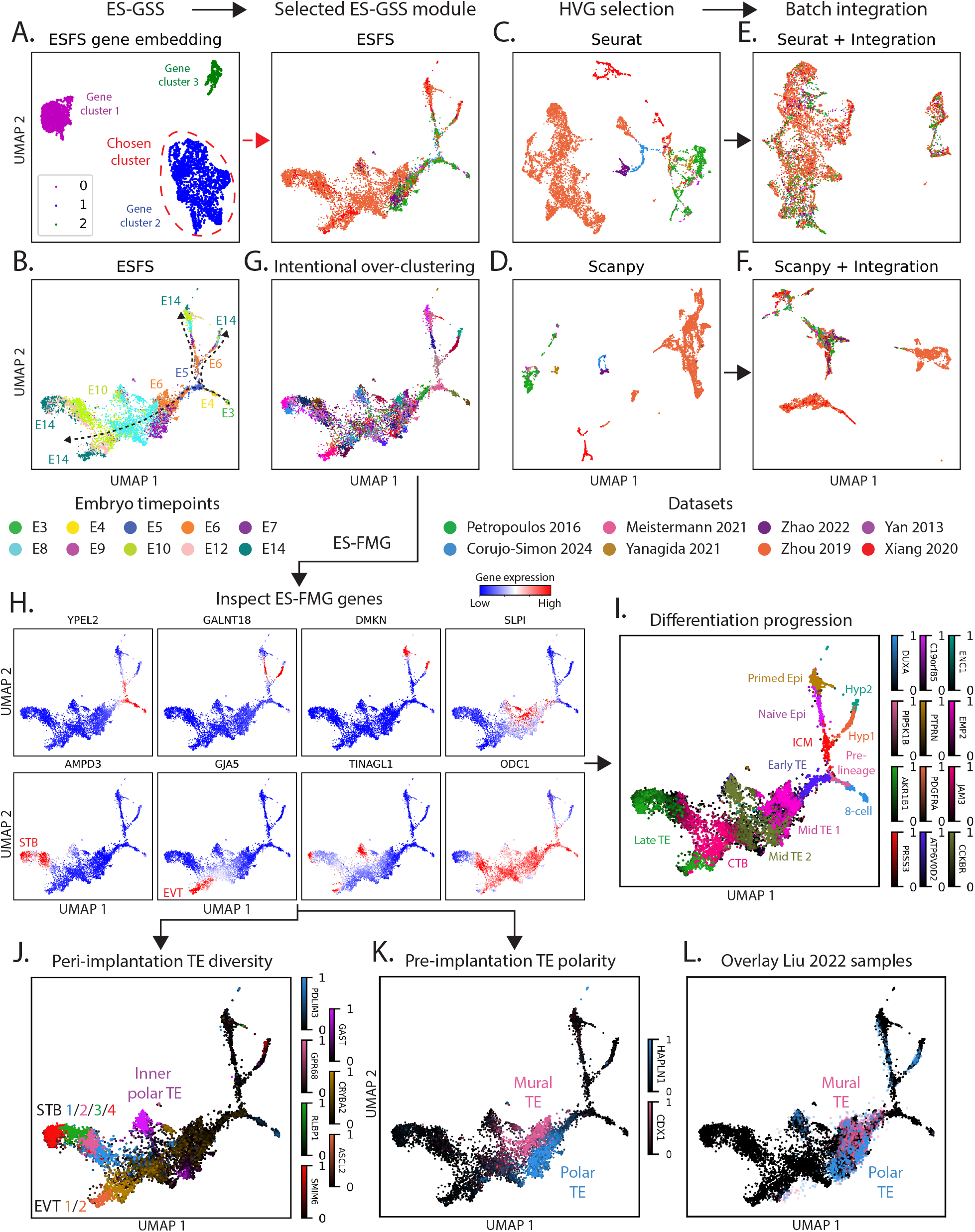
ESFS unbiasedly reveals complex gene expression dynamics in early human embryos. **A**. Subsetting the combined embryo data scRNA-seq counts matrix to 2435 genes identified by ES-GSS (left panel) produces a high resolution E3-E14 human embryo development embedding (right panel) without latent space extraction or batch integration. **B**. Timepoint labels reveal coherent temporal progression across differentiation trajectories. **C**,**D**. Analysing the same data with Seurat or Scanpy HVG selection fails to mitigate interference from dataset batch effects. **E**,**F**. While batch integration from the Seurat and Scanpy workflows performs well at dataset mixing, unintentional data distortion produces embeddings with lower developmental coherency and interpretability. **G**. Intentional over-clustering of samples using the ESFS geneset acts as the input for the ES-FMG algorithm. **H**. A selection of the top 300 marker genes identified by ES-FMG reveals a diverse range of gene expression profiles. **I-K**. ES-FMG reveals waves of sequential expression during early human embryo development (**I**), trophectoderm diversity (**J**) and peri-implantation mural vs. polar TE (**K**). **L**. Unsupervised projection of independent laser dissected mural Vs. polar TE scRNA-seq data reiterates reference data identities.

To compare ESFS-derived embedding with those generated using conventional workflows, we applied Seurat and Scanpy for feature selection followed by batch correction. In contrast to ESFS feature selection, UMAP embeddings constructed using highly variable genes (HVGs) from the Seurat and Scanpy pipelines showed pronounced batch separation, with cells clustering primarily by dataset of origin rather than biological state (Fig 2C,D). We therefore applied dataset integration using Seurat’s *IntegrateData*() function and Scanpy’s implementation of the Harmony algorithm^39^ via *scanpy*.*external*.*pp*.*harmony_integrate*(). Although both approaches improved dataset mixing (Fig 2E,F), the resulting embeddings failed to recover continuous differentiation trajectories or clear lineage bifurcations. This suggests that latent-space integration can blur biologically meaningful neighbourhood relationships, particularly between stable and transitional cellular states.

By contrast, the ES-GSS gene set identifies a subset of features in which biological signal consistently outweighs technical variation, enabling the construction of low-dimensional embeddings that faithfully reflect developmental structure without requiring explicit batch correction. As a result, ESFS preserves cell–cell neighbourhoods across datasets and reveals coherent trajectories and branching events directly from gene expression space. This highlights ESFS as a principled alternative to latent-space integration, offering an interpretable and biologically grounded framework for analysing sparse, heterogeneous scRNA-seq data.

#### ES-FMG highlights molecular diversity in early human embryos

Having generated a high-resolution embedding using the 2435 ES-GSS-selected genes, we then used the same gene set to intentionally over-cluster the dataset in preparation for combinatorial analysis with ES-CCF. Over-clustering was performed using unsupervised Leiden clustering with a relatively high resolution parameter of 8 (Fig 2G). ES-CCF then identified optimised combinations of clusters for each gene, which served as input for the ES-FMG algorithm. We used ES-FMG to select the top 300 genes that most effectively revealed distinct expression patterns across the combinatorial cluster space. By contrast with conventional differential expression analysis, which typically identifies genes enriched within individual clusters, ES-FMG enables the discovery of robust expression profiles spanning one or more clusters, thereby capturing both cluster exclusive and overlapping transcriptional programs (Fig 2H).

Within the top 300 ES-FMG genes, we found genes that captured the emergence of Hypoblast, Epiblast (Epi) and Trophectoderm (TE) populations during early human embryo development (Fig 2I, Fig S3A). For early human Hypo cells, ES-FMG identified *PDGFRA* as an early hypoblast marker proceeded by *ENC1* upregulation. These findings are supported by immunofluorescence analysis of *PDGFRA* and *GATA4* ^16^, with *GATA4* being the highest correlated gene with *ENC1* in our analysis (Fig S3B).

Epiblast can be separated into naïve and primed Epi populations, representing two distinct stages of pluripotency^40^. In our analysis ES-FMG identifies *PTPRN* as a marker gene that captures primed Epi cells, as corroborated by canonical markers *FGF2* and *SFRP2* ^41^ (Fig S3B). Similarly, *LEFTY2* and *KLF17* have been previously identified as genes that distinguish naïve Epi from primed Epi^15^ which is corroborated by our embedding (Fig S3B) and our ES-FMG analysis that more specifically identifies this same distinct Epi population via *C19orf85* (Fig 2I, Fig S3A).

While the inner cell mass (ICM) has been established as a precursor to Epi and Hypo lineages in the mouse^42^, its identification in human embryos has been elusive due to low cell numbers and batch effects. Only recently has an ICM-like population been reported in human scRNA-seq datasets^43^. As such, it is notable that ES-FMG identified *PRSS3* and *LAMA4* as marker genes since these genes have been independently identified as human ICM markers through computational and immunofluorescence analysis, respectively^5^.

In humans, TE cells are essential for blastocyst formation and uterine implantation^44^. Our ES-FMG analysis identified sequential gene expression waves along the TE lineage from E5 to E14 (Fig 2I, Fig S3A). The TE program begins in E5–E6 embryos with *ATP6V0D2*, previously reported as an early TE marker^45^. This is followed by *EMP2*, another ES-FMG identified gene, knockdown of which is known to impair implantation^46^. Subsequently, *CCKBR* expression emerges, which is implicated in preparing the TE for invasion^47^, and precedes the upregulation of *MMP2* and *MMP3* (Fig S3C), consistent with prior findings that *CCKBR* activity induces MMP expression^48^.

ES-FMG also identified *JAM3* as expressed at the bifurcation between syncytiotrophoblast (STB) and extravillous trophoblast (EVT) lineages (Fig 2H,I, Fig S3A), suggesting that *JAM3* may mark the cytotrophoblast (CTB) precursor population. While *JAM3* has not been directly validated as a CTB marker *in vivo, LGR4* (Fig S3D), the sixth most correlated gene with *JAM3*, is a marker of STB/EVT precursor states in *in vitro*^49^. To verify *JAM3* as a CTB marker, we analysed data from a peri-implantation trophoblast differentiation study^50^ and projected it onto our ESFS UMAP embedding (Fig S3E). Encouragingly, in our ESFS embedding at the boundaries between the proposed *JAM* 3^+^ cells and the STB/EVT populations we also observed a boundary between the CTB and STB/EVT cells annotated by West et al. 2019 (Fig S3E, dashed lines). Finally, *AKR1B1* was identified as a marker of late-stage (E12–E14) TE cells, expressed in both STB and EVT lineages, in agreement with earlier reports^51^.

Together, these results demonstrate that the ESFS workflow enables accurate, unbiased, and detailed characterisation of early human embryo scRNA-seq data, even in the presence of limited cell numbers and pronounced batch effects. By identifying marker genes that capture Epi, Hypo and TE early human developmental trajectories without requiring supervision, ES-FMG offers a powerful approach for systematically uncovering both established and previously uncharacterised cellular states during complex developmental processes.

#### Uncovering trophectoderm diversity during human embryo implantation with ES-FMG

After confirming that the ESFS workflow accurately recapitulates previously described gene expression dynamics during early human embryogenesis, we asked whether ES-FMG reveals previously uncharacterised but biologically meaningful cell populations. Prior scRNA-seq studies of early human trophoblast development have primarily relied on discrete clustering, known marker genes, and differential expression analysis to classify cells into cytotrophoblast (CTB), syncytiotrophoblast (STB), and extravillous trophoblast (EVT) states^45,50,52^. While effective at recovering broad lineage identities, this approach lacks the sensitivity to resolve structured heterogeneity within these lineages. By contrast, the objective function of ES-FMG aims to identify a small set of genes that maximally separate biologically meaningful groups of cells from the rest of the dataset.

We identified several genes that stratify TE cells into distinct subpopulations (Fig 2J,K). Along the CTB-to-STB differentiation trajectory, we identified four waves of sequential *PDLIM3* -*GPR68* - *RLBP1* -*SMIM6* gene expression (Fig 2J, Fig S4A). Additionally, ES-FMG highlighted *AMPD3* as a marker gene expressed across the latter three waves of gene expression (Fig S4B), demonstrating the utility of identifying markers that capture continuity across the combinatorial cluster space. Similarly, along the CTB-to-EVT axis, we identified sequential *CRYBA2* -*ASCL2* expression (Fig 2J, Fig S4C), as well as a third gene (*GJA5*) capturing both subpopulations (Fig S4D).

We next focused on a distinct subpopulation of TE cells that ES-FMG identified as having enriched expression of *GAST* (Fig 2J, Fig S4E). This population is largely composed of cells from E6–E10 embryos (Fig 2B). To our knowledge, this population has not been described in prior scRNA-seq analyses of the human embryo. However, recent work by Corujo-Simon et al. 2024 provides compelling evidence for the emergence of a multi-layered polar TE population during implantation. Using sequential fluorescent labelling, they showed that *GATA*3^+^ cells from the E5–E7 mural TE divide and are recruited into internal positions of the polar TE. Visualising *GATA3* expression in our ESFS embedding confirmed that the *GAST* ^+^ population is strongly *GATA*3^+^ (Fig S4E), raising the possibility that this population represents the mural TE-derived internalised cells described by Corujo-Simon et al. 2024.

In our embedding, experimentally validated polar TE markers such as *SDC1* and *PGF* ^45^ are expressed in both STB cells and the *GAST* ^+^ population, supporting the proposed polar identities (Fig S4G). However, these genes show minimal expression in E6–E7 embryos, limiting their utility for distinguishing mural from polar TE at early stages. Among the ES-FMG marker genes, we identified *CDX1* and *HAPLN1* as specifically marking two distinct subpopulations in E6–E7 embryos (Fig 2K, Fig S4H). *CDX1* has previously been suggested as a mural TE marker^45^, while *HAPLN1* has not, to our knowledge, been associated with polar TE.

To assess the validity of *CDX1* and *HAPLN1* as early-stage mural and polar TE markers, respectively, we used an independent scRNA-seq dataset in which human embryos were laser-dissected into polar and mural regions before sequencing^45^. By projecting these samples onto our reference embryo embedding, we assessed similarity to different embryonic cell states. Projection of these samples revealed clear enrichment of mural samples in the *CDX*1^+^ population and polar samples within the *HAPLN* 1^+^ population. As noted in the original study, some label ambiguity is expected due to imperfect boundary selection during laser dissection. Nonetheless, these findings nominate *CDX1* and *HAPLN1* as promising spatial markers of mural and polar TE identity during the peri-implantation window.

In addition, two of the marker genes identified by ES-FMG, *SLPI* and *IRS1*, are specifically upregulated in the identified *GAST* ^+^ population and *CDX*1^+^ mural population (Fig S4I), providing evidence of a shared transcriptional signature.

Taken together, our analysis suggests that the *GAST* ^+^ cells are more transcriptionally related to E6-E7 mural TE than E6-E7 polar TE, but by E9-E10 these cells adopt a more polar molecular identity. These results support the findings of Corujo-Simon et al. 2024, in which E5–E7 mural TE cells separate and are recruited into internal positions of the polar TE.

In summary, the ESFS workflow enables comprehensive characterisation of both exclusive and overlapping Epi, Hypo, and TE cell states, even in the presence of substantial batch effects. Notably, ESFS facilitated the identification of a distinct *GAST* ^+^ TE population that was not detected in previous analyses of these data. Critically, this is achieved without the need for feature extraction or latent space approximation, both of which can introduce computational artefacts that obscure biological signals and complicate downstream tasks. For instance, previous studies have required bespoke, reference-specific pipelines to align query samples into a common latent space before projecting them onto embryo embeddings^34,35^. In contrast, by operating directly in gene expression space, ESFS offers a simple, robust, and generalisable framework for projecting query samples onto high-resolution UMAP embeddings while reducing the risk of artefactual distortions.

### ESFS captures gene expression compartmentalisation in mouse colon spatial transcriptomic data

Spatial transcriptomics (ST) has emerged as a transformative approach for profiling gene expression while preserving tissue architecture^53^. The ability to account for the physical neighbourhoods of cells has led to bespoke software for tasks such as characterising microenvironments and cell-cell communication^54,55^. However, the core scRNA-seq analysis tasks of data pre-processing and identification of robust gene expression patterns remain. Indeed, the biological insights gained from ST depend directly on the quality and resolution of cell state identification.

To test the ability of ESFS to isolate robust, biologically distinct expression states from ST data, we took Visium 10X ST data from the Parigi et al. 2022 study investigating mouse colon spatial composition. The proximal to distal axis of the mouse colon is stratified into 3 distinct layers (Fig 3A). The outer serosa layer covers the muscular wall and faces the peritoneal cavity. The lamina-propria (LP) resides above the serosa between the epithelium and the muscularis mucosae. Finally, intestinal epithelial cells (IEC) are located at the apical border of the gut wall, in direct contact with the food and microbiota present in the lumen.

**Figure 3:**
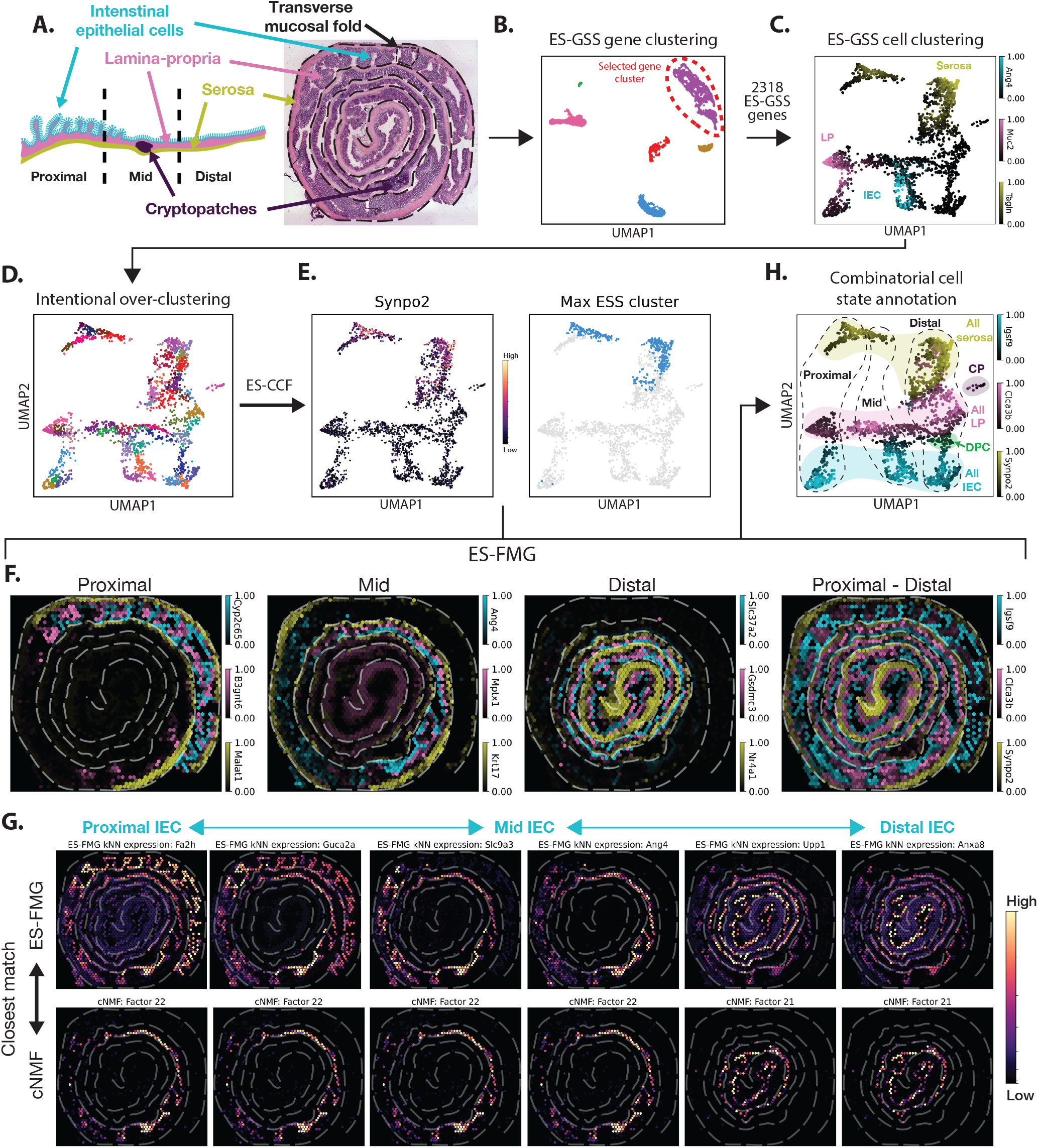
ESFS captures gene expression compartmentalisation in mouse colon spatial transcriptomics. **A**. “Swiss roll” mouse colon spatial transcriptomics data from Parigi et al. 2022 can be broadly characterised by three spatial compartments along the proximal-distal axis. **B**. ES-GSS identifies a set of 2318 genes that produces a high resolution cell state UMAP (**C**.). **D**. Intentional over-clustering of the data serves as the input to ES-CCF to identify groups of cells that maximally correlate with individual genes. **E**. ES-CCF efficiently identifies where the expression of a gene (left panel) is maximally enriched within a population of over-clustered cells (right panel, blue samples). **F**. ES-FMG identifies genes that stratify distinct IEC, LP and Serosa populations within and across proximal, mid and distal compartments. **G**. ES-FMG reveals gene expression gradients within IEC samples along the proximal to distal axis that failed to be identified by cNMF.

Despite applying current best practices, including non-negative matrix factorisation (NMF), Parigi et al. 2022 were unable to distinguish LP from IEC towards the distal end of the colon. This discrepancy—between anatomically distinct compartments and their indistinct transcriptional signatures in the ST data—highlights a key limitation of current methods and provides an opportunity to evaluate whether ESFS can improve upon prior analyses. Here, we demonstrate that ESFS not only resolves this limitation, but also achieves higher-resolution identification of complex cell states and uncovers biologically meaningful gene expression gradients that remain obscured by feature extraction approaches such as NMF.

To ensure a fair comparison, we used the spatial transcriptomics (ST) scRNA-seq counts matrix from Parigi et al. 2022, comprising 2,604 sequencing spots that had undergone initial quality control (e.g., removal of low-quality spots). We filtered out genes expressed in fewer than 30 spots, retaining 14,402 genes for downstream feature selection using ES-GSS. ES-GSS identified a set of 2318 genes (Fig 3B), which when used to subset the scRNA-seq counts matrix, produced a highly structured UMAP of cell similarity, without requiring PCA-based dimensionality reduction (Fig 3C).

To annotate cell neighbourhoods, we used validated marker genes from Parigi et al. 2022 for the serosa (*Tagln*), lamina propria (LP; *Muc2*), and intestinal epithelial cells (IEC; *Ang4*) (Fig 3C). Spatial visualization of these markers revealed expression in distinct sub-populations along the proximal-distal axis, consistent with the UMAP clustering (Fig S5A). *Tagln, Muc2* and *Ang4* showed enrichment in the mid, proximal and distal compartments respectively.

Before applying ES-CCF, we intentionally over-clustered the data using Leiden clustering with resolution parameter set to 8, generating 55 discrete clusters (Fig 3D). ES-CCF was then applied to identify coarse-grained, noise-resistant groupings of cells by maximising the Entropy Sort Score (ESS) between each gene and its optimally correlated combination of Leiden clusters (Fig 3E). The resulting ESS cluster-gene associations were passed to ES-FMG and the top 200 genes that best captured distinct expression signatures across the combinatorial cluster space identified.

#### ESFS identifies Serosa-LP-IEC stratification along the entire proximal-distal axis

Parigi et al. 2022 successfully identified spatially distinct gene expression programs demarcating the proximal, mid, and distal colon, but clear stratification of serosa, LP, and IEC populations was only observed in the proximal region. In the mid region, only IEC-related populations were identified. In the distal region, only IEC and serosa layers were recovered, LP was absent.

Using the expert-curated analysis from Parigi et al. 2022 as a high-quality reference, we compared the performance of our ESFS pipeline to that of cNMF^25^ (Experimental Procedures).

By inspecting the 200 genes selected by ES-FMG we were able to identify marker genes that capture Serosa-LP-IEC stratification along the entire proximal-distal axis. These display clear structure in both the spatial coordinate space (yellow-pink-blue, Fig 3F) and UMAP space (Fig S5B). To compare these results with those from cNMF, we took each of the expression profiles identified by ES-FMG and used the cosine similarity metric to find their most similar cNMF factor (Fig S6). In agreement with Parigi et al. 2022, cNMF accurately identifies factors that capture distinct Serosa, LP and IEC populations in the proximal compartment (Fig S5C). However, while Parigi et al. 2022‘s analysis could only identify a factor relating to the IEC in the mid domain, cNMF identifies factors relating to mid IEC and LP (Fig S5C). Despite this increased resolution, only ES-FMG identified gene expression profiles enriched in mid spatial compartment for all 3 of the Serosa-LP-IEC populations (Fig 3F, Fig S5C). Similarly, while ES-FMG convincingly revealed Serosa-LP-IEC layering throughout the distal region (Fig 3F), cNMF failed to capture this spatial organisation (Fig S5C). Strikingly, whereas neither Parigi et al. 2022 nor cNMF recovered factors that span Serosa–LP–IEC layering across the full length of the colon (Fig S5C), ES-FMG identified gene expression signatures that were enriched in each of the 3 cell compartments across the entire proximal/distal axis (Fig 3F).

Together, these results demonstrate that ESFS matches or outperforms NMF in the unsupervised identification of biologically meaningful gene expression patterns in spatial transcriptomics data. Notably, ESFS enables combinatorial cell state annotation, automatically uncovering hierarchical structures within cell states (Fig 3H)—a clear advantage over cNMF and conventional discrete clustering approaches.

#### ES-FMG and cNMF identify known and new gene expression profiles

Having characterised Serosa-LP-IEC stratification along the proximal-distal axis, we investigated other gene expression profiles identified by ES-FMG and cNMF. During their analysis, Parigi et al. 2022 identified and verified lymphoid structures by factors enriched with B-cell associated genes. For example, an NMF factor defined the colonic LP comprising the expression of genes characteristic of plasma cells such as *Igha, Jchain, Igkc*. Consistent with this, ES-FMG identified *Igha* as one of the top 200 marker gene candidates, producing a spatial expression pattern matching the factor identified by Parigi et al. 2022 and factor 51 identified by cNMF (Fig S5D). Cryptopatches (CPs) and isolated lymphoid follicles (ILF) are small, lymphoid cell aggregates found in the intestinal LP that play a role in gut immunity. ES-FMG and cNMF identified expression signatures that match the CP and ILF populations identified by Parigi et al. 2022 (Fig S5E, F). ES-FMG also identified *Elavl4* as a top ranked marker gene, a gene that is associated with the enteric nervous system (ENS)^57^. Inspection of *Elavl4* expression profile and the most similar cNMF factor show correspondence with the ENS factor characterised by Parigi et al. 2022 (Fig S5G). These results confirm that ES-FMG and cNMF capture the diverse range of gene expression signatures characterised by Parigi et al. 2022.

In addition to expression signatures identified by Parigi et al. 2022, we asked if ES-FMG identified other spatially restricted gene expression profiles. Transverse mucosal folds (TMFs) are anatomical structures in the proximal region of the mouse colon, with the stem cell niche/crypt in the trough of the fold closest to the serosa, and mature differentiated cells at the tip (Fig 3A). By inspecting the top marker genes identified by ES-FMG we found genes expressed specifically in the proximal compartment of the colon. The expression of these was restricted to either the tip (*Car1*, Fig S5H) or the base (*Prkg2*, Fig S5I) of the proximal TMFs. The identification of *Car1* as a marker for the tip of proximal colon TMFs agrees with previous observations^58^. Alkaline intestinal phosphatase (*Alpi*) is expressed in differentiated enterocytes but not in immature crypts^59^. *Alpi* was also identified by ES-FMG as a significant gene, and inspection of *Alpi* shows that its gene expression profile matches closely to that of *Car1* (Fig S5J), further validating their co-expression at the tip of the TMFs. Conversely, the proximal zone of mouse colons of *Prkg2* -/- mice exhibit crypt hyperplasia (an increase in the length and/or number of crypts), suggesting it plays an antiproliferative and prodifferentiation role^60^. These results are consistent with our findings placing *Prkg2* expression at the base of the TMFs (Fig 3L).

These results confirm that ES-FMG identifies cell state and spatially restricted gene expression profiles in a computationally unbiased manner. By matching the *Car1* and *Prkg2* expression profiles identified by ES-FMG with their most similar cNMF factors, we can confirm that cNMF also identifies a wide range of distinct biological signatures. However, a key distinction between the ESFS workflow and cNMF is that ES allows analysis in gene expression space rather than a latent space. In the next section we demonstrate that this difference allows the identification of heterogeneous expression profiles missed by NMF approaches.

#### ES-FMG reveals gene expression heterogeneity obscured by cNMF

Dimensionality reduction techniques such NMF are commonly applied to scRNA-seq data to summarise gene co-expression patterns into a smaller set of latent factors. This compression often yields interpretable features that highlight dominant expression profiles. However, by projecting data into a reduced latent space, these methods risk distorting biological signals or obscuring subtle gene expression gradients of interest^6^. In contrast, the objective function of the ES-FMG algorithm directly identifies a set of *N* genes whose expression profiles maximally correlate with minimally overlapping cell groupings (Materials and Methods). By doing so, ES-FMG isolates genes that capture distinct expression structures, including those that span transitional states between dominant profiles. We therefore hypothesised that ES-FMG-selected features would better preserve gene expression gradients than latent features extracted by cNMF.

Within the 200 genes identified by ES-FMG, we identified a set of 6 genes expressed in a gradient in IEC cells along the proximal to distal axis (Fig 3G). Taking the expression profiles of the 6 genes and using cosine similarity to identify the most similar cNMF factors, we find that cNMF has compressed the identified expression gradient into 2 latent factors (Fig 3G). Hence these results indicate that by simplifying the ST data into a set of latent factors, cNMF has unintentionally obscured the identification of biologically relevant gene expression gradients.

Conversely, by identifying an optimal set of potential marker genes directly from the gene expression space, ES-FMG produces highly interpretable results that can simplify the identification of subtle but distinct gene expression profiles in scRNA-seq data. From a computational modelling perspective these results suggest that the genesets identified by ES-FMG more faithfully elucidate molecular signatures that bridge the gaps between the predominant cell states within a dataset than the latent factors derived from cNMF.

Taken together, our analysis shows that ESFS facilitates comprehensive characterisation of the mouse colon ST data, identifying local, global and graded expression signatures that are missed by current best practices.

### ESFS captures spatial heterogeneity within and between glioblastoma tumours

Glioblastoma (GBM) is an aggressive primary brain tumour characterised by diffuse infiltration and marked molecular heterogeneity. Recent studies have applied spatial transcriptomics (ST) to glioblastoma samples to investigate tumour architecture and microenvironments^61–65^. However, the highly unstructured and variable organisation of glioblastomas within and across tumours makes these data particularly challenging to interpret. Previous approaches have relied on integration strategies to extract broad expression programs, often at the cost of masking tumour-specific heterogeneity^61^, or required manual, sample-by-sample annotation to resolve local expression microenvironments^62^. These challenges present an opportunity to test whether ESFS enables the analysis of multiple samples in a single workflow directly from the gene expression space.

In the following we show that the ESFS distils complex, heterogeneous tumour data into interpretable representations of cellular states. The ability of ES-FMG to recover combinatorial expression patterns directly from the gene expression space simplifies the characterisation of shared and tumour specific spatial heterogeneity in glioblastoma, and offers a route for investigating other tumour types.

#### ESFS facilitates simultaneous analysis of multiple tumours

To analyse inter-tumour heterogeneity and the complex spatial organisation of GBM tumours Ravi et al. 2022 developed a bespoke, supervised dataset integration pipeline to identify gene expression programs that distinguished broad cell states across tumour samples. While this approach successfully recovered genesets reflecting distinct GBM cell states, it remained difficult to pinpoint marker genes for spatially discrete cellular niches both within and between tumours. In follow-up work, Kueckelhaus et al. 2024 highlighted that conventional differential gene expression analysis is unreliable in predicting gene expression with spatial dependencies. As such, the authors introduced a ST analysis tool that allows users to manually define linear or radial gradients within tumour sections and search for spatially restricted gene expression profiles along these axes. Although effective in specific scenarios, this approach struggles to generalise across heterogeneous tumours and may miss spatial niches that lack clear anatomical axes.

Motivated by these challenges, we asked whether the ESFS workflow could provide a robust and generalisable framework for identifying spatially distinct gene expression profiles in glioblastoma. To do so, we compiled 15 glioblastoma ST samples from Ravi et al. 2022 (Fig 4A) into a single dataset and applied the ESFS workflow. Using ES-GSS, we identified a set of 2194 genes (Fig 4B), and found that this geneset produced a UMAP embedding with clearly resolved sub-populations (Fig 4C). We estimated copy number variation (CNV) scores (*infercnvpy* ^66^) as a proxy for malignant cells in each ST spot and found that malignant samples were concentrated toward the centre of dense regions of the embedding, indicating that the ESFS projection primarily organises samples by malignant states (Fig S7A). We also observed a distinct group of spots with consistently low malignancy scores. We suspected these might represent low-quality samples and confirmed this by inspecting the total transcript abundance per spot (Fig S7B).

**Figure 4:**
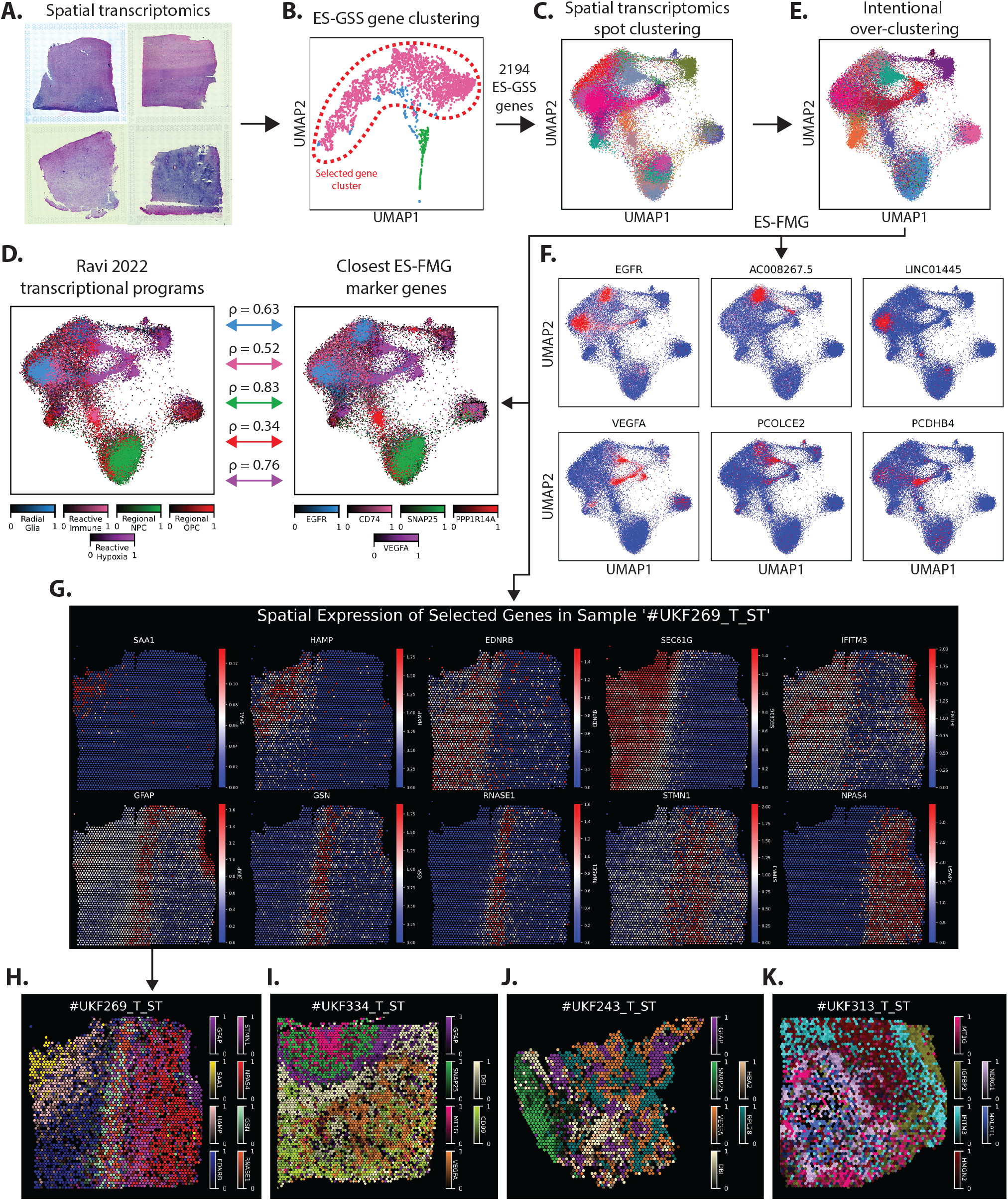
ESFS reveals shared and tumour-specific expression patterns in glioblastoma spatial transcriptomics (ST) data. **A**. Representative examples of glioblastoma ST samples included in the analysis. **B**. ES-GSS identifies a cluster of 2194 genes that generates a high-resolution, low-dimensional embedding. **C**. UMAP embedding of ST spots based on the ES-GSS gene cluster. Each colour corresponds to one of the 15 glioblastoma samples included in the combined dataset. **D**. Intentional over-clustering was applied using Leiden clustering to prepare the data for ES-FMG analysis. **E**. Comparison of canonical glioblastoma transcriptional programs from Ravi et al. 2022 (left) with the activity of the five most correlated genes among the top 200 marker genes identified by ES-FMG (right). Double-headed arrows indicate Pearson correlation values between each transcriptional program and ES-FMG gene activity. **F**. ES-FMG identifies marker genes enriched in both discrete clusters and across overlapping subpopulations. **G**. Spatial maps of individual tumours showing spatially restricted expression of selected ES-FMG genes. **H-K**. ES-FMG enables robust detection of spatially restricted gene expression patterns across heterogeneous tumours with varying topologies and levels of complexity.

Importantly, we were able to generate this embedding without processing each sample individually or applying computational integration methods that risk artificially removing biologically meaningful heterogeneity between tumours.

#### ES-FMG identifies canonical glioblastoma expression signatures

To investigate the biological significance of the separate ST spot populations in the ESFS UMAP embedding, we first asked whether they recapitulated previously characterised GBM expression programs. In Ravi et al. 2022, the authors defined five distinct transcriptional programs — radial glia, reactive immune, regional neural progenitor cells (NPCs), regional oligodendrocyte progenitor cells (OPCs), and reactive hypoxia. Using the genesets associated with each program, we scored each ST spot using Scanpy’s *score_genes()* function (Fig 4D). This revealed clusters of spots enriched for the radial glia, regional NPC, and reactive hypoxia programs.

We intentionally over-clustered the data via Leiden clustering (Fig 4E) and applied ES-FMG to identify an optimised set of 200 marker genes capturing robust expression profiles within and between tumours. To assess the overlap with the five programs identified by Ravi et al. 2022 we identified the gene from the 200-marker ES-FMG set with the highest Pearson correlation with each program. Four out of five of these genes, *EGFR, CD74, SNAP25* and *VEGFA* were also found in GBM transcriptional program genesets Ravi et al. 2022. For the radial glia, regional NPC and reactive hypoxia programs, we found a strongly correlated marker (*ρ >* 0.6), indicating that ESFS recovered transcriptionally similar patterns in an unsupervised manner (Fig 4D). For the reactive immune program, we observe overlap with the radial glia program (Fig 4D). Here, the top ES-FMG marker was *CD74*, with *ρ* = 0.52. This is consistent with previous work describing the reactive immune program as a hybrid state that spans multiple GBM sub-populations^61,67^. Moreover, Zeiner et al. 2015 found that *CD74* expression is largely confined to microglia/macrophages in gliomas, supporting the notion that this program reflects the tumour microenvironment rather than tumour-intrinsic states.

Finally, we found that the regional OPC program did not localise to a specific population in the embedding, and the top-correlated ES-FMG gene (*PPP1R14A*) had only moderate similarity (*ρ* = 0.34). However, recent findings from Tang et al. 2024 identified *PPP1R14A* as a glioma oligodendrocyte marker gene, suggesting that ES-FMG may have recovered an OPC- or OPC-like population.

Together, these results demonstrate that ESFS recovers known GBM transcriptional programs in an unsupervised and generalisable manner. Moreover, the ESFS embedding indicates that canonical signatures such as the radial glia and reactive hypoxia programs may exist across multiple distinct spot populations (Fig 4D), indicating there may be additional structured heterogeneity within transcriptional programs.

### ESFS highlights shared and tumour specific expression profiles without supervision

Encouraged by ESFS’s ability to identify broad glioblastoma expression programs, we next investigated whether the ESFS workflow could highlight gene expression profiles found in distinct subsets of tumours and tumour microenvironments.

We began by asking whether within the 200 ES-FMG geneset we could find genes that appeared to partition the broad transcriptional programs characterised by Ravi et al. 2022 (Fig 4D). Indeed, we were able to identify genes such as *EGFR, AC008267*.*5* and *LINC01445* that are upregulated in different sections of the radial glia program (Fig 4F). A similar pattern can be observed for the reactive hypoxia program when inspecting *VEGFA, PCOLCE2* and *PCDHB4* (Fig 4F). To verify the unsupervised marker gene analysis from ES-FMG, we performed a complementary supervised analysis using Leiden clustering with default parameters (Resolution = 1), and generated ranked gene lists for each of the resulting clusters (Fig S7C-E). As expected, ES-FMG correctly identified marker genes with expression profiles that matched top ranked genes identified through broad Leiden clustering (Fig S7E,F). For example, *EGFR, AC008267*.*5, LINC01445, VEGFA, PCOLCE2* and *PCDHB4* were ranked 4, 3, 3, 8, 2 and 1 respectively in their closest broad Leiden cluster. These results demonstrate that ES-FMG not only identifies broad GBM transcriptional programs but also highlights distinct sub-populations without requiring iterative clustering supervision.

Having shown that the ESFS workflow can unbiasedly resolve complex heterogeneity in the gene expression space, we next asked whether ES-FMG could simplify the identification of marker genes for spatial niches in glioblastoma within a chaotic, multi-tumour dataset. To illustrate the difficulty of this task, we examined the spatial expression of *VEGFA, PCOLCE2*, and *PCDHB4* across four tumours (Fig S7G), and found that while each gene exhibited spatial restriction in some tumours, they lacked consistent patterns across samples. Similarly, comparing the activity of the radial glia program in a single tumour against top-ranked radial glia genes reveals that relying on broad transcriptional programs can unintentionally obscure spatially restricted expression of interest (Fig S7H). Indeed, inspecting the tumour sample composition of the broad Leiden clusters shows that expression states are not evenly distributed across tumours (Fig S7D). This variability highlights the limitations of relying on discrete clustering or broad expression signatures to identify spatially restricted marker genes across tumours.

We hypothesised that because ES-FMG selects candidate marker genes directly from the gene expression space – rather than from a latent representation – we can be confident that the spatial markers reflect true gene expression signals, rather than smoothed or imputed profiles potentially inherited from neighbouring tumour contexts.

Starting with a single sample, we inspected the ST expression profiles of the 200 genes selected by ES-FMG and found genes with clear spatially restricted expression patterns within a single tumour (Fig 4G, H). Repeating this process across additional tumours, we identified genesets that act as markers of distinct microenvironments in individual tumour samples (Fig 4I–J, Fig S8A–C).

Amongst the spatially distinct genes identified via ES-FMG, several have been previously linked to human GBMs, such as *GFAP* ^65,70^, *NDRG1* ^71^, *MALAT1* ^72^, and *HAMP* ^73^. *GFAP* has been shown to mark astrocytes and reactive tumour-associated astrocytes at the invasive edge of GBMs. These findings are corroborated by our identification of *GFAP* at an invasive edge in tumours #UKF269_T_ST and #UKF334_T_ST (Fig 4G–I, Fig S8A). However, in tumour #UKF243_T_ST, *GFAP* expression appears to form distinct islands rather than a continuous boundary (Fig 4J, Fig S8B), while in #UKF313_T_ST, *GFAP* is notably absent (Fig S8D). Conversely, in tumour #UKF313_T_ST, *NDRG1* forms a circular boundary around a *MALAT1* -positive population (Fig 4K, Fig S8C), but shows no discernible boundary in tumours #UKF269_T_ST and #UKF334_T_ST (Fig S8D). In tumour #UKF243_T_ST, *NDRG1* forms islands of expression similar to, but independent from, those marked by *GFAP* (Fig S8D). Together these results indicate that tumour #UKF243_T_ST is an example of GBM where chaotic spatial organisation means that computational methods that search for linear or radial boundaries would struggle to identify spatially restricted marker genes. Finally, although *HAMP* expression has previously been reported as elevated in GBMs, among the four tumours visualised, we observed spatially distinct *HAMP* expression only in tumour #UKF269_T_ST, with sparse and heterogeneous expression in the remaining three tumours.

More generally, while a given geneset might reveal distinct and meaningful spatial expression within one tumour, applying the same set to other tumours often leads to a deterioration in the ability to identify spatial niches (Fig S8D). These results confirm the heterogeneous nature of GBMs. Moreover, visualising these spatially restricted genesets in UMAP space (Fig S8E) underscores the importance of ES-FMG’s ability to explore the combinatorial expression space. For example, genes such as *GFAP* show spatially restricted expression in multiple tumours, yet their expression profiles span disparate regions of the UMAP embedding limiting their identification through a single set of discrete clusters.

Taken together, these results demonstrate ESFS’s ability to process GBM expression data and reveal global, shared, and tumour-specific expression programs within a single, easy-to-use workflow. Notably, whereas recent state-of-the-art methods for glioblastoma analysis often rely on supervised annotation or assumptions of linear or radial anatomical structure, ESFS instead focuses downstream analysis and tumour characterisation on a relatively small set of informative genes, without requiring prior knowledge or iterative semi-supervised clustering. As a result, the ability of ESFS to robustly and unbiasedly identify spatially restricted expression profiles represents a powerful tool for the detailed characterisation of glioblastoma and other heterogeneous systems.

### Characterisation of spatial, temporal and neurogenesis expression dynamics of spinal cord development

In vertebrate embryos, neural tube (NT) development is governed by three overlapping regulatory programs: spatial and temporal patterning, and neuronal differentiation^26,74,75^ (Fig 5A–B). Spatial patterning specifies distinct cell types based on their position along the dorsal-ventral axis. Temporal patterning determines cell identity according to the developmental timing of progenitor differentiation. The neuronal differentiation program controls the transition from proliferative progenitor to post-mitotic neuron. The combinatorial nature of gene expression patterns across these three axes presents a major challenge for scRNA-seq analysis of the NT. In mouse datasets, conventional workflows such as Seurat and Scanpy struggled to resolve clear differentiation trajectories across spatial domains without relying on domain expertise, and do not comprehensively capture molecular signatures associated with the temporal or neurogenesis programs^26,76^.

**Figure 5:**
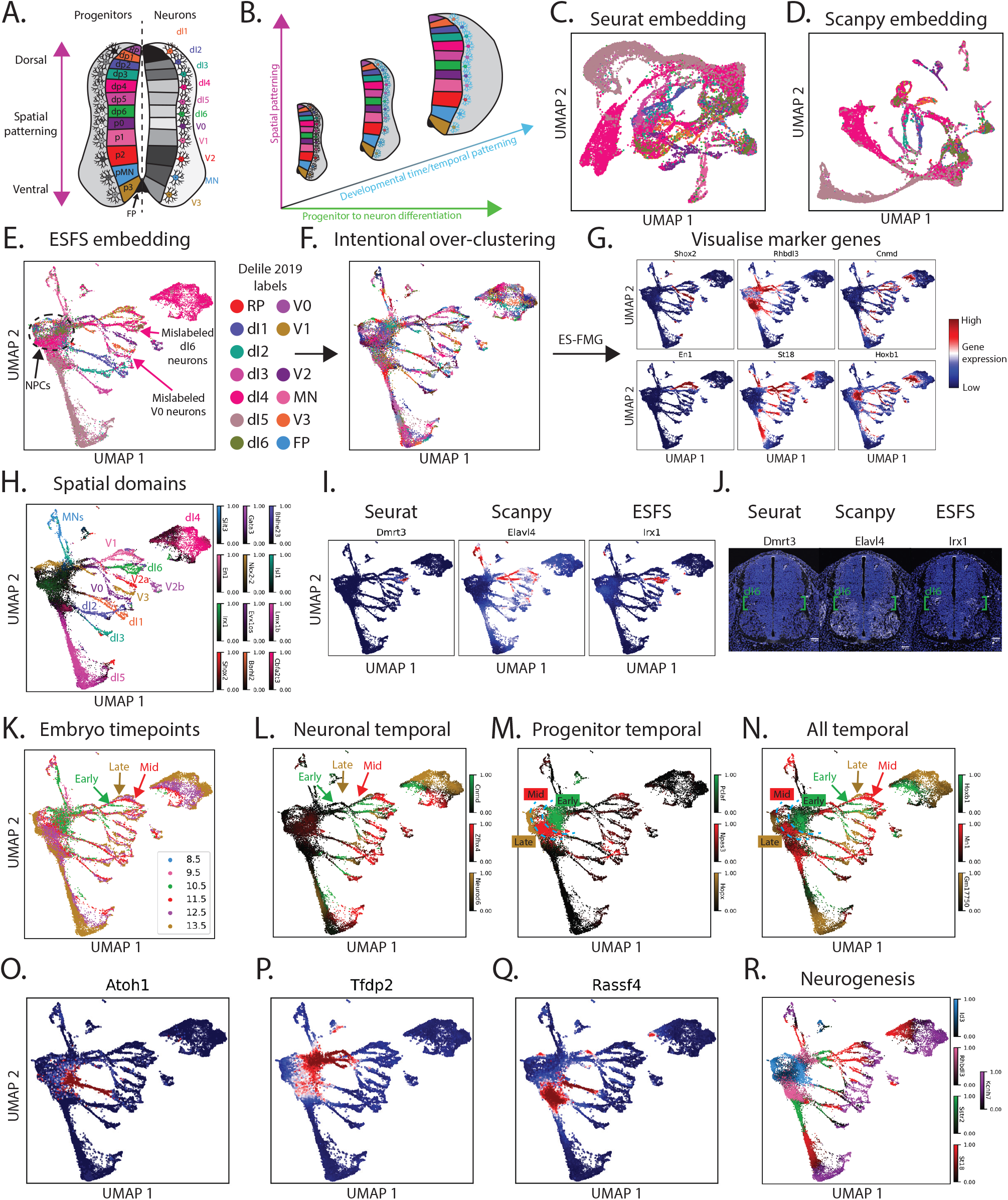
ESFS unbiasedly captures the spatial, temporal and neurogenesis expression dynamics of neural tube development. **A**,**B**. NT differentiation is regulated by at least 3 distinct programmes. 1) Spatial patterning along the dorsal-ventral axis (A.) generates spatially distinct neuronal populations. 2) Progenitors that initiate differentiation at different time points (B., black-blue neurons) produce molecularly distinct neurons within spatial domains. 3) Progenitor-neuron progression may be tracked by markers of neuronal maturity. **C**,**D**. Conventional Seurat and Scanpy scRNA-seq analysis workflows fail identify coherent progenitor to neuron cell states. Panels C, D and E share the same legend. **E**. ESFS reveals distinct neuron populations with clear progenitor-neuron trajectories. **F**. Intentional over-clustering of the data prior to application of our ES-FMG algorithm. **G**. Visualisation of some of the top marker genes identified by ES-FMG. **H**. ES-FMG identifies marker genes capturing neurons from different spatial domains. **I**. Top 2 dI6 marker genes identified by ESFS are more robust than those identified by Seurat or Scanpy. **J**. HCR staining of mouse E11.5 NT confirms superior robustness of ESFS dI6 marker genes. **K**. Embryonic time points from which NT scRNA-seq data was derived. **L-N**. ES-FMG reveals temporally restricted gene expression profiles in neuron (L.), progenitor (M.) and neuron + progenitor (N.) populations. **O-Q**. ES-FMG identifies genes highlighting intermediate dI1 interneurons (O.), intermediate ventral interneurons (P.) and intermediate dI1-dI5 interneurons (Q.). **R**. ES-FMG identifies genes that broadly display waves of sequential expression as NPCs differentiate into interneurons.

Here, we reanalyse the scRNA-seq dataset from Delile et al. 2019 and show that ESFS enables high-resolution reconstruction of neural tube development. ESFS reveals gene expression profiles that distinctly stratify cells according to their spatial position, developmental timing, and differentiation maturity. By recovering both established and previously unannotated cellular states, without requiring prior labels or supervision, ESFS effectively decomposes complex developmental scRNA-seq datasets into biologically interpretable and functionally relevant subpopulations.

#### ESFS reveals high-resolution neural tube differentiation dynamics

Consistent with prior analyses Delile et al. 2019, Gupta et al. 2024, applying the HVG methods in Seurat and Scanpy to the Delile et al. 2019 scRNA-seq dataset resulted in embeddings that clearly separated dI4 and dI5 populations (Fig 5C,D). However, visual inspection of the remaining interneuron subtypes shows they either cluster together indistinctly or fail to form coherent trajectories along the neurogenesis axis. These limitations complicate downstream clustering and pseudotime analysis. Indeed, both Delile et al. 2019 and Gupta et al. 2024 relied on manual annotation using known marker genes, rather than unsupervised clustering or enrichment analysis—limiting their capacity to uncover novel cell states and regulatory programs.

Using ESFS’s ES-GSS algorithm, we identified a set of 835 genes that produced UMAP embeddings with smooth, continuous neurogenesis trajectories linking progenitors to interneuron populations across spatial domains (Fig 5E). We then applied ES-FMG to select the top 400 genes that capture distinct expression profiles (Fig 5F,G). From this set, we identified 26 genes that highlight complex spatial, temporal, and neurogenesis related expression dynamics underlying neural tube development (Fig S9), which we discuss in detail below.

These results demonstrate the ability of the ESFS workflow to generate highly interpretable and biologically structured representations from complex scRNA-seq datasets. By removing the reliance on prior knowledge or assumptions, ESFS allows the identification of new, biologically relevant expression dynamics.

#### Spatial domains

In the developing neural tube (NT), *Bmp* and *Shh* signalling along the dorsal–ventral axis allocate neural progenitor cells (NPCs) into distinct spatial domains (Fig 5A, Fig S9A), which subsequently differentiate into specific neuronal subtypes. In previous analyses^26,76^, unsupervised clustering failed to resolve molecularly distinct interneuron populations. Consequently, cell-type annotation relied on supervised identification using experimentally validated marker genes. While effective for confirming known populations, this approach risks biasing analyses toward predefined identities, potentially overlooking rare or previously uncharacterised cell types. For example, although the authors attempted to annotate dI6 neurons using canonical markers such as *Dmrt3*, they identified only a small number of matching cells.

Inspection of the ESFS embedding revealed that the small group of dI6 neurons identified in Delile et al. 2019 is nested within a larger population previously labelled as dI4 neurons (Fig 5E). Similarly, the previously identified V0 neuron population is surrounded by samples annotated as dI4 neurons. Motivated by these observations, we asked whether, within the top 400 marker genes identified by ES-FMG, we could recover specific markers for each of the 12 interneuron populations, without using prior cell-type labels.

Among the 400 genes selected by ES-FMG, we identified clear and specific marker genes for 11 of the 12 spatial domains (Fig 5H, Fig S9A). Although we did not find a gene uniquely marking the dI3 population, ES-FMG identified *Isl1* as a top-ranked gene expressed in both the dI3 and motor neuron (MN) populations, consistent with previous findings^77^. Many of the domain-specific markers identified by ES-FMG have prior experimental support, including *En1, Nkx2-2, Gata3* ^78,79^, and *Barhl2* ^80^. Additionally, because each marker gene identified by ES-FMG is associated with a specific cell cluster (Materials and Methods), we can use the ranked gene list for each ES-FMG marker gene to identify co-expressed genes within the same spatial domain. For instance, although we found no prior literature validating *Evx1os*, the second-highest ranked gene correlated with the *Evx1os* ES-FMG associated cluster is *Evx1* (Table S1) — a well-established marker of V0 interneurons^81^.

Motivated by ESFS’s ability to identify marker genes for distinct spatial domains without supervision — and noting that the canonical dI6 marker gene, *Dmrt3*, labels only a subset of dI6 neurons — we asked whether the top-ranked ESFS gene for the putative dI6 population, *Irx1* (Fig 5H,I), might serve as a more robust marker. We further sought to compare the performance of ESFS against marker gene identification methods from Seurat and Scanpy. As reported previously^26,82^, Seurat ranked *Dmrt3* highest for the dI6 population, while Scanpy identified *Elavl4*, which appeared broadly upregulated across multiple interneuron populations. Targeted hybridisation chain reaction (HCR) staining of E11.5 mouse neural tube sections validated these observations (Fig 5J), showing minimal *Dmrt3* expression, broad non-specific upregulation of *Elavl4*, and strikingly specific *Irx1* expression within the dI6 domain. These results indicate that ESFS more accurately prioritises population-specific gene expression than conventional differential expression methods.

Together, these results demonstrate that ESFS accurately identifies spatially resolved gene expression programs and interneuron populations within the NT. By facilitating unbiased data characterisation that does not rely on prior knowledge or manual curation, ESFS allows the identification of novel, biologically relevant gene expression profiles.

#### Temporal patterning

In parallel to spatial patterning, a temporal regulatory program operates in the neural tube (NT) that systematically changes progenitor identity over developmental time, such that progenitors located in the same spatial domain generate molecularly distinct neuronal populations depending on when they initiate differentiation^74,83–85^. This temporal program progressively shifts the molecular identity of progenitors. Specifically, Sagner et al. 2021 and Zhang et al. 2025 reported that early-born neurons express *Onecut1* /*Onecut2*, intermediate-born neurons express *Zfhx3*, and late-born neurons express *Nfia*/*Nfib*. The temporal program is also apparent in NPCs, with early progenitors expressing *Lin28a*/*Lin28b* and *Nr6a1*, followed by *Npas3* and *Sox9* in intermediate stages, and finally *Nfia*/*Nfib* in late-stage progenitors.

In the high-resolution ESFS embedding, we observed that while the dominant separation of cells reflects progenitor and interneuron populations across spatial domains, an additional stratification is visible based on embryonic time point (Fig 5K). When we visualised the temporal marker genes identified by Sagner et al. 2021 and Zhang et al. 2025 on the ESFS embedding, their expression patterns recapitulated temporal dynamics in both progenitor and neuronal populations (Fig S10). For instance, *Lin28a* and *Nfia* expression coincided with early (E10.5) and late (E12.5–13.5) NPCs, respectively (Fig S10A). These findings indicate that feature selection through ES-GSS can unbiasedly improve data interpretability such that spatial and temporal populations become distinct in a single embedding.

This prompted us to search for genes within the top 400 ES-FMG markers that stratify progenitor and neuronal populations according to developmental time. We first identified early-, mid-, and late-expressed genes within neurons (Fig 5L). Notably, although *Onecut1* /*Onecut2* appeared biased toward early ventral neurons, ES-FMG identified *Cnmd* as an early neuron marker with broader representation across both ventral and dorsal domains (Fig 5L, Fig S9B). At the same time, ES-FMG captured the ventral *Onecut2* signature as biologically meaningful, as the ES-FMG gene *D930028M14Rik* (within the top 400 genes) lists *Onecut2* as its third most correlated gene (Fig S10B). Similarly, *Zfhx4*, an ES-FMG-selected gene, marked intermediate-born neurons (Fig 5L), with Zhang et al. 2025’s *Zfhx3* being the second most correlated gene in the *Zfhx4* cluster (Fig S10B). Late-born neuron markers such as *Nfia* also appeared in the top 400 ES-FMG genes, although *Nfia* expression extended beyond the interneuron subpopulations into mid- and late-stage NPCs (Fig S10B). Conversely, the ES-FMG gene *Neurod6* showed relatively specific expression in late interneurons and negligible expression in NPCs (Fig 5L, Fig S9B). Hence, ES-FMG facilitated unbiased identification of genes relating to more neuron specific temporal identities than previous supervised investigations.

Next, we investigate whether ES-FMG could identify temporal patterning within the progenitor population. In the Delile et al. 2019 dataset, we found that no combination of the temporal genes reported by Zhang et al. 2025 could effectively stratify early, mid, and late NPCs without also labelling a substantial portion of interneurons (Fig S10A,C). However, within the ES-FMG gene set we identified *Pclaf* and *Hoxp* as early and late progenitor markers respectively, with minimal expression in interneurons (Fig 5M). For the mid-progenitor population, we could not easily identify a specific marker visually, so we queried which of the 400 ES-FMG gene clusters was most correlated with *Npas3*, a known mid-temporal progenitor gene^85^. Encouragingly, *Npas3* ranked second within the ES-FMG cluster defined by *Foxp2* (Fig S9B), suggesting that although *Foxp2* was not an obvious mid-progenitor marker by inspection, ES-FMG successfully identified a strong quantitative signal for this intermediate temporal state.

Finally, within the ES-FMG geneset, we identified genes marking temporal expression programs across both NPC and interneuron populations (Fig 5N, Fig S9B). To our knowledge, this global pattern has not previously been reported, suggesting a previously unrecognised temporal signature spanning the entire neural tube differentiation hierarchy. For a ranked gene list of other genes following these novel gene expression profiles see Table S1.

Together, these results show that ESFS not only recovers known markers of temporal patterning, but also reveals new gene candidates for stratifying temporally patterned NPCs and interneurons in the developing NT. Moreover, the observation that ES-FMG selected genes exhibit greater mutual exclusivity than those identified by standard experimental/statistical workflows provides an example of the usefulness of the optimisation objective driving the ES-FMG algorithm (see Materials and Methods).

#### Neurogenesis

Neurogenesis is characterised by the transition of progenitors into neurons. Intermediate cell states spanning the progenitor to neuron transition remain poorly defined. To better understand the regulatory dynamics of NT neurogenesis, we identified marker genes that distinguished differentiating cells from both NPCs and mature neurons, within and across spatial domains.

Gupta et al. 2024 compared scRNA-seq data from *in vitro* dorsal interneuron differentiation protocols and *in vivo* mouse NT datasets to identify genes marking the progenitor-to-neuron transition. This enabled the identification of markers for differentiating progenitors across dorsal domains, for example, *Tfap2b* expression in intermediate dI2–5 interneurons (Fig S10D). However, because the analysis was limited to dorsal interneuron development, some genes proposed as dorsal-specific were also found to be expressed ventrally. For instance, while *Pax2* has been proposed as a marker of dI4 and dI6 interneurons, it is also known to be expressed in V1 interneurons — a pattern corroborated by our analysis (Fig S10E). Similarly, while *Gsg1l* was proposed as a specific marker of dI1 differentiation, ESFS revealed its expression also spans ventral neuronal differentiation (Fig S10E).

To demonstrate the value of an unsupervised marker gene identification, we searched the top 400 ES-FMG genes for marker genes of intermediate progenitor-interneuron populations. *Atoh1* was identified as a more specific marker of dI1 differentiation than *Gsg1l* (Fig 5O). In addition to recovering *Tfap2b*, ES-FMG highlighted *Rassf4*, which marks differentiating progenitors across dI1–5 domains, and *Tfdp2*, which captures progenitor-to-neuron transitions in ventral regions (Fig 5Q,R). These findings illustrate the utility of ESFS in generating interpretable, high-resolution representations of complex datasets by unbiasedly detecting combinatorial expression patterns underlying neurogenesis.

A complementary study by Yu et al. 2025 combined human and mouse scRNA-seq with spatial transcriptomics (ST) to track gene expression and cell migration during NPC-to-interneuron differentiation. Because their dataset included cells from all spatial domains, they were able to identify markers of differentiating NPCs expressed across spatial domains, such as *Sstr2* and *St18* (Fig S10F). However, to define intermediate states, Yu et al. 2025 partitioned their data into three discrete clusters (progenitors, precursors, and neurons) followed by differential expression analysis. This coarse-grained, supervised approach limits resolution, making it difficult to capture continuous gene expression dynamics beyond the boundaries of the three predefined states. For example, while *Sstr2* and *St18* were designated as precursor markers, our ESFS embedding reveals that their expression is largely sequential rather than concurrent (Fig S10E).

To explore this further, we used the top 400 ES-FMG genes to identify five genes exhibiting sequential expression from progenitors to interneurons across both dorsal and ventral spatial domains (Fig 5P, Fig S9C). The high-resolution, combinatorial search space of the ES-FMG algorithm confirmed five genes, including both *Sstr2* and *St18* ^86^, while also revealing additional markers that resolve finer transitions during neurogenesis. Collectively, these genes offer a more precise panel for staging the maturity of differentiating progenitors in both *in vivo* and *in vitro* settings.

In conclusion, the complexity of NT differentiation, with distinct spatial, temporal and neurogenesis regulatory programmes, presents a challenging dataset for scRNA-seq analysis workflows. Without use of prior knowledge or predefined marker genes, ESFS disentangles the Delile et al. 2019 dataset into highly interpretable gene expression signatures supported by both literature and experimental validation. Notably, ESFS identifies temporal and neurogenesis expression profiles that exist across multiple transcriptionally distinct spatial domains. This capability arises from ES-CCF and ES-FMG providing a tractable framework for searching combinatorial cluster space, a task that remains computationally intractable for conventional marker gene algorithms.

## 3 Discussion

In this study, we introduce Entropy Sorting Feature Selection (ESFS), a machine learning workflow for analysing high-dimensional transcriptomics data. Conventional high-throughput transcriptomic analysis pipelines typically rely on dimensionality reduction and batch integration to de-noise datasets - approaches that risk introducing computational artefacts which can obscure biologically meaningful signals. By contrast, ESFS uses the mathematical framework of Entropy Sorting to identify sets of genes marking biologically relevant cell states and trajectories. By demonstrating that high-resolution embeddings can be achieved directly from gene expression space, our work challenges the prevailing assumption that single cell transcriptomic data necessitates denoising through latent representations or data imputation.

Once biologically informative expression heterogeneity is revealed by ES-GSS, ES-CCF and ES-FMG can be applied to identify genes with distinct expression profiles. Crucially, ES-CCF and ES-FMG enable users to search for an optimal set of features that capture robust expression profiles across the combinatorial cluster space, thereby circumventing the conventional need to iteratively define a single sub-optimal clustering resolution. As a result, these tools provide an efficient and unbiased framework for uncovering complex hierarchical gene expression patterns that may be missed by methods reliant on rigid clustering or predefined lineage structures. Moreover, these tools enable the derivation of minimal genesets that capture meaningful molecular diversity within heterogeneous cell populations, streamlining downstream analysis and experimental design.

For all four datasets presented in this study, ESFS improved the interpretability of high-dimensional transcriptomics data by uncovering complex expression states that were not detected using conventional workflows. The approach demonstrated robustness to common confounders, including batch effects and variation in sequencing depth. In developmental datasets, such as the human embryo and mouse neural tube, ES-GSS revealed fine-grained, biologically coherent developmental trajectories that were obscured by other approaches. In spatial transcriptomics datasets from the mouse colon and human glioblastoma, ES-FMG enabled the unsupervised identification of spatially restricted gene expression patterns without requiring prior annotation. In the mouse colon, ESFS recovered local and global Serosa–LP–IEC stratification signatures that were not resolved by NMF-based methods. In glioblastoma, ESFS revealed shared and tumour-specific cellular states and microenvironments by identifying robust expression programs directly from gene expression space, avoiding distortions introduced by latent-space transformations.

Together, these findings underscore the potential of Entropy Sorting–based methods to distil complex biological data into an interpretable gene expression space, with minimal introduction of computational artefacts. By alleviating the need for data compression, ESFS offers a more faithful foundation for building advanced machine learning and AI tools that can exploit subtle but biologically meaningful expression heterogeneity. Further, we anticipate that applying ESFS to even higher-dimensional datasets – such as chromatin accessibility or multi-omics data – will yield comparable, if not greater, improvements in data interpretability. More broadly, Entropy Sorting provides a generalisable framework for uncovering significant feature relationships and identifying outlier data points without parametric assumptions - paving the way for more transparent, robust, and biologically grounded data analysis and mathematical modelling across a diverse range fields.

## 4 Experimental Procedures

### Data and code availability

Instructions for installing the ESFS software and example workflows for reproducing results from this manuscript are available at the following GitHub repository - https://github.com/aradley/ESFS. Data objects for reproducing results may be found at the following Figshare repository - https://figshare.com/s/4e445e7fa03cc4ccd289

Raw data for all single cell and spatial transcriptomics data are publicly available and maybe found in the following locations:

Mouse colon spatial transcriptomics - Gene expression omnibus, GSE169749.

Human embryo single cell RNA sequencing count matrices were retrieved from the public repository provided by Zhao et al. 2025 (https://petropoulos-lanner-labs.clintec.ki.se/dataset.download.html).

The mouse neural tube single cell RNA sequencing counts matrix may be found in the public repository provided by Delile et al. 2019 (https://github.com/juliendelile/MouseSpinalCordAtlas).

Glioblastoma spatial transcriptomics data was retrieved from the public repository provided by Ravi et al. 2022 (https://datadryad.org/dataset/doi:10.5061/dryad.h70rxwdmj).

### ESFS parameter selection

For each of the datasets analysed in this manuscript, ESFS was applied to improve data interpretability via the ES-GSS algorithm, followed by identification of robust cell state marker genes using the ES-CCF and ES-FMG algorithms. When using ES-GSS, users must select the number of top-ranked genes to carry forward for downstream analysis. As with highly variable gene selection, choosing how many genes to include is often an iterative process. Likewise, for ES-FMG, users may tune the number of distinct marker genes to identify, as well as a resolution parameter that controls the degree to which selected marker gene expression profiles are allowed to overlap.

Below, we describe the parameter choices of the ESFS workflow used to generate the presented results, alongside sensitivity analyses for the ES-GSS top-ranked gene selection and ES-FMG parameters. For Python Jupyter notebooks containing the code used to apply ESFS to each dataset in this manuscript, see: https://github.com/aradley/ESFS/tree/main/Example_Workflows.

#### ES-GSS top ranked gene selection

For each dataset presented in this manuscript, ES-GSS was used to identify a set of genes that, when used to subset the corresponding scRNA-seq or spatial transcriptomics count matrix, significantly enhanced our ability to resolve distinct cell populations and gene expression dynamics. To facilitate this, ES-GSS provides a suite of functions for gene importance weighting, gene set clustering, and gene set selection. The primary user-defined parameter is the number of top-ranked genes to carry forward after gene importance weighting. In panels A–C of Fig S12, Fig S11, Fig S13 and Fig S14, we visualise the effect of varying this parameter for the mouse colon, human embryo, mouse neural tube, and human glioblastoma datasets, respectively.

In panels A, we show how changing the number of top-ranked genes affects the qualitative structure of UMAP embeddings generated using the most informative gene set - typically the one that best distinguishes canonical cell types or reveals biologically meaningful developmental trajectories. For datasets such as the mouse colon and glioblastoma spatial transcriptomics, varying the number of top-ranked genes had only a minor effect on UMAP structure. In contrast, both the human embryo and mouse neural tube datasets showed clear improvements in the resolution of cell groupings and developmental trajectories at the final gene set sizes selected by ES-GSS.

In panels B, we highlight the importance of gene set clustering and set selection. Focusing on the chosen top-ranked genes for each dataset, we compare UMAP embeddings generated using either all selected genes or individual genesets. These comparisons reveal substantial variability in the biological structures captured by different sets. As with the number of top-ranked genes, the impact of gene set selection is dataset-dependent. For example, in the mouse colon data, using all top-ranked genes yields a UMAP embedding similar to that produced by gene set 5 (Fig S12B). In contrast, in the embryo and neural tube datasets, selecting a set aligned with a relevant developmental structure significantly improves data interpretability (Fig S11B, Fig S14B).

Finally, in panels C, we assess the biological relevance of each gene set via gene ontology (GO) enrichment analysis. Across datasets, sets identified as capturing biologically meaningful structure in UMAP space were consistently enriched for relevant GO terms. For instance, in the mouse colon dataset, gene set 5 is enriched for terms related to protein catabolism and biosynthesis—processes consistent with gut physiology (Fig S12C). Other genes sets were enriched for GO terms less specific to gut or colon identity.

#### ES-CCF and ES-FMG top marker gene selection

Following ES-GSS gene selection, the resulting cell-cell neighbourhoods can be used as an input for the ES-CCF and ES-FMG algorithms. As described in the ESFS workflow section (Materials and Methods), the main input to ES-CCF is a set of cell cluster labels from an intentionally over-clustered transcriptomics dataset (Fig 1iii). The choice of clustering method and resolution is user-defined. For the mouse colon, human embryo and human glioblastoma datasets, we used Leiden clustering with resolution parameters of 9, 8 and 3, respectively. For the mouse neural tube data, we found that hierarchical clustering with 250 target clusters more effectively separated subpopulations within spatial domains. Once over-clustering is complete, ES-CCF is applied to compute a per-gene maximum ESS cluster matrix (Fig 1vii), which is then passed to ES-FMG.

ES-FMG requires two user-defined parameters: the number of marker genes to identify (*N*) and a resolution parameter (*ρ*) that controls the allowable overlap between marker gene expression profiles. The *N* parameter is conceptually similar to specifying the number of latent features in dimensionality reduction approaches such as PCA, NMF, or autoencoders. However, unlike those methods, ES-FMG does not compress the data into a latent space. Instead, *N* determines the number of genes to be selected that best capture robust and distinct expression profiles within the dataset.

The *ρ* parameter governs the stringency of redundancy penalization during marker gene selection (Fig 1ix, equation (18)). Values closer to 1 impose stronger penalties for overlapping expression patterns across marker genes, favouring distinct profiles; values closer to 0 allow more overlap. For instance, in the mouse colon spatial transcriptomics dataset, we set *ρ* = 0.7 to highlight biologically meaningful expression gradients that could be missed by dimensionality reduction methods such as NMF (Fig 3G). In contrast, for the early human embryo, mouse neural tube and human glioblastoma data, we set *ρ* = 1 to prioritise the identification of clearly distinct cell states.

To assess the robustness of ES-FMG to different choices of *N* and *ρ*, we performed grid searches across a wide range of values for each dataset. For benchmarking, we used the sets of biologically meaningful genes identified in the main figures as ground truth - 19, 21, 26 and 20 genes for the mouse colon, human embryo, mouse neural tube and human glioblastoma datasets, respectively. At each parameter combination, we computed the cosine similarity between these reference genes and their most similar counterparts in the ES-FMG output. Results are shown in panels D of Fig S12, Fig S11, Fig S13, and Fig S14. These analyses demonstrate that ES-FMG is highly robust to both *N* and *ρ*, with cosine similarity values quickly plateauing near 1 as *N* increases.

### Gene expression plots

All cell UMAP embeddings were generated using a nearest neighbour size of 50, a minimum distance of 0.1, and a correlation-based distance metric. To improve interpretability, unless otherwise stated, all gene expression plots for the mouse colon, human embryo, and mouse neural tube datasets (scRNA-seq and spatial transcriptomics) display the *k*-nearest neighbour (kNN) mean expression per data point, with *k* = 50.

For the human glioblastoma dataset, expression heterogeneity across tumour samples is a key focus. Smoothing gene expression across tumours would obscure biologically meaningful micro-environment differences. Therefore, all gene expression plots for glioblastoma display raw expression values without smoothing.

For multi-gene expression plots - either on UMAPs or spatial transcriptomics grids - we applied the following procedure to highlight regions of high activity. First, the expression values for each gene were clipped at the 97.5th percentile. These values were then converted to percentiles, such that, for example, a value at the 80th percentile was mapped to 0.8. For each data point, we selected the gene with the highest percentile value (i.e., the most active gene at that location) to determine the displayed expression level. Each gene was assigned a unique colour and corresponding colour bar. Finally, plotting order was determined by percentile rank, such that lower percentile values were drawn first and higher percentile values last, ensuring that high-expression regions remained visually prominent.

### Consensus non-negative matrix factorisation (cNMF) comparison

To benchmark the performance of ES-FMG in identifying distinct gene expression profiles, we compared its results to those generated by the widely used feature extraction method, consensus non-negative matrix factorisation (cNMF)^25^, using the mouse colon spatial transcriptomics dataset. We used the spatial organisation of the dataset to evaluate how well each method could recover biologically interpretable, spatially restricted expression patterns.

Following the recommended cNMF workflow, we began by conducting a model stability analysis, running cNMF with the number of latent features (factors) set between 20 and 200 in increments of 10. We fixed the number of input genes to 2318, matching the number of genes selected by the ES-GSS algorithm to ensure a fair comparison. The upper limit of 200 latent factors was chosen to match the number of marker genes selected by ES-FMG. Stability analysis indicated a peak in factor reproducibility when the number of factors was set to 90, which we used for downstream analyses.

To compare the outputs of the two approaches, we computed the cosine distance between each of the 90 cNMF latent features and each of the 200 ES-FMG marker gene kNN-features. We then visualised representative gene expression profiles from each method side by side (Fig 3G). This allowed us to assess where ES-FMG and cNMF identified similar or divergent spatial expression patterns, highlighting the strengths and limitations of each approach in capturing biologically meaningful structure in the data.

### Seurat and Scanpy analyses

When comparing outputs from our ESFS workflow with those of the Seurat and Scanpy scRNA-seq analysis workflows, Seurat/Scanpy were used with recommended parameters on identical scRNA-seq counts matrices. Our findings that Seurat and Scanpy workflows struggle to resolve known cell types in the early human embryo and mouse neural tube scRNA-seq datasets is corroborated by previous studies^26,28,34,35,87^.

### *In situ* hybridization chain reactions

To validate dI6 interneuron markers in the mouse neural tube, *in situ* hybridization chain reactions (HCR) were performed on neural tube sections for top ranked genes identified by the Seurat, Scanpy and ESFS workflows. HCR v3.0 probe sets targeting *Dmrt3, Wt1os, Irx1, Irx2, Nrxn3* and *Elavl4* were designed using a custom script (https://github.com/ctucl/pybridizer, Suppermpool et al. 2025) and ordered as pooled DNA oligonucleotides (oPools, Integrated DNA Technologies). HCR amplifiers and all hybridization and wash buffers were obtained from Molecular Instruments. E11.5 mouse embryos were collected in PBS, fixed overnight in 4% paraformaldehyde at 4°C, rinsed in PBS and cryoprotected in 15% sucrose in phosphate buffer overnight at 4°C. Embryos were embedded in a gelatin and sucrose solution (7.5% gelatin, 15% sucrose in phosphate buffer) and snap-frozen in isopentane chilled on dry ice. Transverse neural tube sections (14 m) were prepared on a Leica CM3050S cryostat and mounted on Superfrost Plus slides (Thermo Scientific, 10149870). Slides were stored at -80°C until further processing. HCR was performed following the Molecular Instruments HCR v3.0 protocol for tissue sections. Slides were post-fixed in 4% PFA for 15 minutes, step-wise rehydrated in 25% increments of PBST (0.1% Tween 20 in PBS; Sigma-Aldrich, P2282) on ice, treated with protease K (10 g ml^−1^; VWR International, A3830.0500) for 10 minutes at 37°C and briefly post-fixed again in PFA (20 minutes). Sections were equilibrated in 5× SSCT (SSC buffer with 0.1% Tween 20; Invitrogen, 15557-044), pre-hybridized for 30 minutes at 37°C with probe hybridization buffer (Molecular Instruments) and incubated overnight at 37°C with probes at a final concentration of 24 nM. After washing in 5× SSCT, amplification was performed by incubating slides in amplification buffer for 5 minutes, then overnight at room temperature in the dark with hairpins. The hairpin solution was prepared by snap cooling the hairpins for 30 minutes and then mixing them to obtain a 60 nM solution in amplification buffer. Sections were washed in 5× SSCT, counterstained with DAPI and mounted in ProLong Glass Antifade Mountant. Images were acquired using a Leica SP8 scanning confocal microscope.

## Supporting information

Table S1 - Mouse neural tube marker genes

## 5 Acknowledgements

We thank Ariel Levine, Brian Roome, Joao Silva, Dimitris Volteras, Elena Corujo-Simon, Lawrence Bates, Oliver Stegle and Magnus Rattray for critically reading and constructive discussions; the Francis Crick Institute Software Engineering & AI STP for software development support; Florian Hubl for advising on the interpretation of the mouse colon spatial transcriptomics data; Afnan Azizi for help with HCR probe design.

## 7 Author contributions statement

Conceptualisation: A.R., J.B.; Data curation: A.R.; Formal Analysis: A.R., G.B.; Funding acquisition: J.B.; Investigation: A.R.; Methodology: A.R., C.S.; Project administration: J.B.; Resources: J.B., G.B.; Software: A.R., C.S.; Supervision: J.B.; Validation: A.R., G.B.; Visualisation: A.R.; Writing – original draft: A.R.; Writing – review & editing: A.R., J.B, G.B., R.P-C.

## Materials and Methods

### Entropy Sorting theory updates

ESFS uses the Entropy Sorting (ES) mathematical framework to non-parametrically calculate correlation and significance scores between genes in scRNA-seq data. These scores—termed the Entropy Sort Score (ESS) and Error Potential (EP), respectively—quantify the strength and reliability of gene co-expression patterns across single-cell datasets. Our initial formulation of the ES framework, developed for discretised data, was introduced in Radley et al. 2023, and subsequently extended in Radley, Boeing, and Smith 2024 to support continuous data in a computationally efficient manner. Unlike alternative information-theoretic approaches such as mutual information and transfer entropy, ES avoids the computational overhead associated with estimating multivariate joint probability density functions. However, the original derivation for discretised data imposed implicit constraints that limited the flexibility of the ES framework in downstream applications. Here, we introduce theoretical updates that lift these constraints, improving the generalisability of ES to more complex datasets. These refinements enhance the framework’s ability to detect significant relationships between features (in transcriptomics data each gene is a feature), and thereby increase the analytical power of tools built on ES such as the ESFS workflow.

#### Introduction to Entropy Sorting nomenclature

In the following we provide an introduction to Entropy Sorting (ES) nomenclature, alongside a nomenclature table (Table S2) and visual representations of key terms and their usage in Fig. S1.

As outlined in Radley et al. 2023, the mathematical framework of Entropy Sorting (ES) stems from a reformulation of the conventional probabilistic conditional entropy (CE) equation – which can be written in the nomenclature of ES (Fig. S1A, Table S2) – into a sorting problem. By inspecting the overlap between non-zero observations in each feature (e.g. gene), we may quantify the degree to which two features correlate with one another.

For any pair of features with values normalized between 0 and 1, it is intuitive that a high overlap of non-zero values reflects strong absolute correlation (e.g., overlap of yellow samples to the right of ESE1 in Fig. S1C,2.). However, strong correlations can also arise in the form of anti-correlation, where the overlap between non-zero values is low (e.g., overlap of yellow samples to the right of ESE2 in Fig. S1C,2.). Here lies an incongruency where identifying the correct methodology for quantifying correlation between two features according to the overlap of non-zero values is dependent on prevalence of zero and non-zero values in each feature. An overview of identifying the correct methodology is provided in Fig. S1C, and discussed in detail later.

To address this incongruency problem, we implement a more general formulation where for any given feature, the non-zero values in individual observations/samples are designated as either minority state values or majority state values as follows:

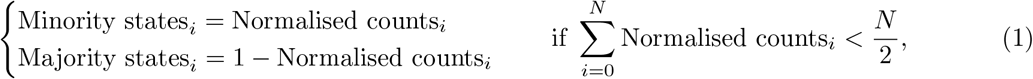

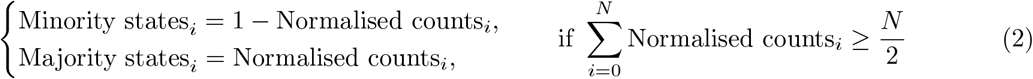

where *N* is the total number of samples, and *i* indexes each individual sample from 1 to *N*.

In the context of gene expression, this formulation interprets “minority states” as referring to whichever of the two – active (non-zero expression) or inactive (zero expression) – occurs to a lesser degree across the *N* observations. Conversely, “majority states” refer to the more prevalent of these two expression states within the same set of observations.

Having identified whether “Normalised counts_*i*_” or “1−Normalised counts_*i*_” capture the minority and majority states respectively, for each of the *N* samples comprising a feature (*F*), we designate the total minority states for feature *F* with subscript *m* such that:

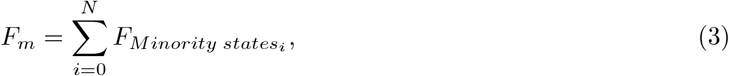

and the total majority states for feature *F* with subscript *M* such that;

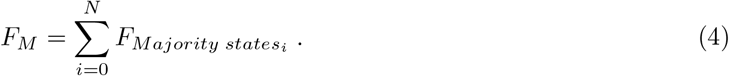

Next, as previously outlined in Radley et al. 2023, for any pair of features being compared against one another, one must be designated the Reference Feature (RF) and the other the Query Feature (QF). This designation is required so that we may identify the correct procedure for calculating ES derived metrics (Fig. S1C, 1.), and is carried out according to the Maximum Entropy Principle such that:

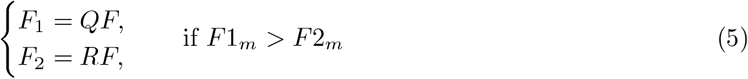

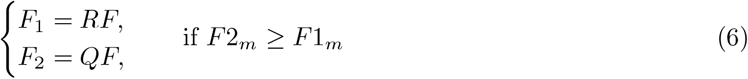

It follows that, *RF*_*m*_ and *RF*_*M*_ refer to total presence of minority or majority states in the Reference Feature (RF) respectively, such that *RF*_*m*_ +*RF*_*M*_ = *N* and *QF*_*m*_ +*QF*_*M*_ = *N*. Graphical representations of these definitions are provided in Fig S1C.

In the probabilistic form of CE (Fig. S1A), *RF*_*m*_, *RF*_*M*_, *QF*_*m*_, and *QF*_*M*_ exist as scalar constants defined by a given pair of features. In addition to these constants, we find 4 dependent scalar variables, *QF*_*m*_, *RF*_*m*_, *QF*_*M*_, *RF*_*m*_, *QF*_*m*_, *RF*_*M*_ and *QF*_*M*_, *RF*_*M*_, representing the co-occurrence of minority and/or majority states between any Query Feature (QF) and Reference Feature (RF) pair (further details below). By substituting any three of these four variables with equivalent terms containing the remaining variable and appropriate constants, we can create 4 new equations containing a single variable that describe the conditional entropy between two features (Fig. S1B). For example, in Fig. S1B, ESE1 represents a rearranged form of the conditional entropy equation, where the sole remaining variable, *RF*_*m*_, *QF*_*m*_, indicates the overlap of minority states between the RF and QF.

Together ESE’s 1-4 and the definitions above form the foundations of the ES framework (Fig. S1C), allowing the quantification of dependent relationships between features through the lens of a sorting problem.

### Limitations of previous derivations of the Entropy Sort Equation

Having outlined fundamental ES notation, here we describe limitations of ES frameworks implemented in previous works^5,15^. ESE1 (Fig S1B) reformulates conditional Shannon entropy in terms of a single variable: the co-occurrence of minority state observations between two features, denoted by the *RF*_*m*_, *QF*_*m*_ scalar. Algorithmically, this overlap is computed as the element-wise sum of minority-state co-occurrences between a RF and QF pair. Since the minimum of two observations is equal to the intersection/overlap of their values, *RF*_*m*_, *QF*_*m*_ may be calculated as:

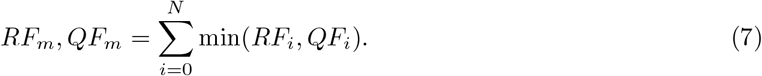

However, the calculation of co-occurrence using equation (7) is only valid when both features being compared have their minority and majority states assigned according to equation (1). If one or both features instead have their minority and majority state assignments determined by equation (2), the interpretation of co-occurrence becomes incongruent. In such cases, the variable *RF*_*m*_, *QF*_*m*_ no longer reflects overlap between equivalent (i.e., like-for-like) observations.

For example, in either of the ESE4 feature pair cases shown in Fig S1C, 2., both *RF*_*m*_ and *QF*_*m*_ are assigned based on the complement of normalised values (i.e., 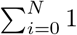 - Normalised counts_*i*_). Attempting to apply equation (7) in this scenario fails, because the minority states correspond to zero-valued observations, resulting in *RF*_*m*_, *QF*_*m*_ = 0. This leads to a spuriously low estimate of the conditional entropy, effectively misrepresenting the relationship between the two features.

In our previous work, we addressed this incongruence by enforcing the use of equation (1) for all features. This was achieved by transforming feature observations such that the sum of each feature’s normalised non-zero values was always less than half the number of samples. Formally, for each gene:

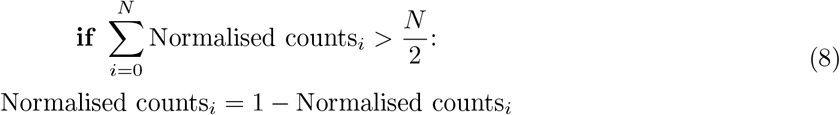

This approach allowed us to circumvent inconsistency in the interpretation of co-occurrence and successfully apply ES in earlier scRNA-seq analysis tools^5,15^. However, as we show in subsequent sections, this constraint introduces limitations – particularly in detecting certain classes of anti-correlated relationships between features.

To overcome these shortcomings, we now generalise the ES framework by incorporating all four Entropy Sort Equation (ESE) variants (Fig S1B). This enables direct quantification of Entropy Sort Scores (ESSs) and Error Potentials (EPs) between feature pairs without requiring transformation of the data to conform to the constraints of ESE1.

#### Using all 4 Entropy Sort Equations to identify previously missed anti-correlations

In Fig S1B, we present the Entropy Sort Equation used in our previous works (ESE1)^5,15^, alongside three additional re-arrangements of conditional Shannon entropy that yield single-variable formulations (ESE2–4). Fig S1C provides a flow diagram outlining how to apply each ESE equation appropriately and compute the Entropy Sort Score (ESS) and Error Potential (EP) metrics for any given pair of features. To facilitate comparisons, we designate one feature as the fixed feature (FF) and the other as the secondary feature (SF). This convention is particularly useful in high-dimensional settings like scRNA-seq, where a single feature (e.g., a gene of interest) is compared systematically against all others. The flow diagram in Fig S1C can then be followed to determine the appropriate ESE equation for each FF/SF pair.

As noted earlier, our previous software tools^5,15^ applied the transformation in equation (8) to force all features into a form compatible with ESE1. For instance, feature pairs conforming to ESE2–4 were transformed such that they could be evaluated with ESE1. When applied to binary data with two discrete states, this transformation has no effect on the ESS or EP, apart from reversing the sign of the ESS in the case of anti-correlated feature pairs. We visualise this in Fig S2A–B, where an ESE2 compatible FF/SF pair (panel A) is transformed into an ESE1 compatible form (panel B), by inverting the expression values of the FF. As expected, the ESS and EP remain consistent (with an opposite ESS sign), indicating correct identification of correlation or anti-correlation.

However, when working with continuous data, this forced transformation can obscure true anti-correlated relationships. As shown in Fig S2C, applying ESE1 to a strong but imperfectly anti-correlated FF/SF pair with non-discrete values can result in a low ESS, incorrectly suggesting a weak relationship. This is because ESE1 quantifies the overlap of like-for-like minority state observations (*RF*_*m*_, *QF*_*m*_ co-expression), rather than the more appropriate alignment of inverse expression. Instead, our flow diagram for ESE usage (Fig 8C) indicates that after inverting the FF (i.e., setting *FF*_*i*_ = 1 − *FF*_*i*_), we may correctly reveal a stronger (larger *ESS*) and significant (*EP >* 0) anti-correlation via the application of ESE2 (Fig S2D).

This example highlights that ESE equations form complementary pairs dependent on the composition of the observed random variables, and selecting the correct equation requires examining both the original FF/SF pair and their suitable inverse form.

In summary, the mathematically rigorous procedure for calculating *ESS* and *EP* between any FF/SF pair involves computing metrics for both the original and the inverted FF (i.e., *FF* and 1 − FF), using the appropriate ESE in each case as per Fig S1C. The *ESS* and *EP* from the scenario yielding the largest absolute *ESS* value may then retained as the least likely arrangment to have occurred by random chance. This updated framework has been implemented in the ESFS software presented in this manuscript, allowing accurate detection of both correlated and anti-correlated relationships in continuous high-dimensional data.

#### Accounting for both divergent and overhang error potentials

The Error Potential (EP) is a significance metric derived from the Entropy Sorting (ES) framework. Analogous to a statistical *p*-value, it quantifies whether an observed relationship between two features is likely to reflect a meaningful dependency or is merely the result of random variation. Specifically, the EP estimates the likelihood that inconsistent observations – those that deviate from the expected conditional relationship – arise from system noise disrupting a true dependency, rather than from a chance correlation between two independent features.

In our previous work^5,15^, we formalised the EP as follows:

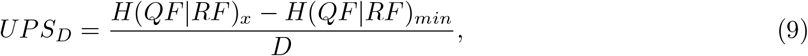

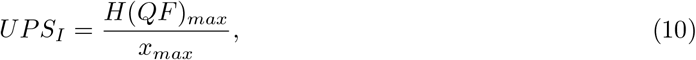

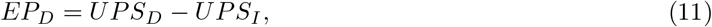

Here, *UPS* stands for uncertainty per sample, where the _*D*_ and _*I*_ subscripts stand for Divergence and Independence respectively. *UPS*_*D*_ and *UPS*_*I*_ represent the uncertainty per sample under the assumptions of feature dependence and independence, respectively. *H*(*QF*|*RF*)_*x*_ is the conditional entropy (CE) at the observed overlap value *x*, which are both calculated according the algorithmic flow diagram in Fig S1C. *H*(*QF*|*RF*)_*min*_ is the minimum possible CE for the RF/QF pair (Fig S2, black arrows). *H*(*QF*)_*max*_ is the maximum entropy (equivalent two perfect independence between the QF and RF), and *D* is the number of divergent samples – those whose observations are inconsistent with the hypothesis of an optimally dependent relationship.

As illustrated in Fig S2A, the values *H*(*QF*)_*max*_, *H*(*QF*|*RF*)_*min*_, and *H*(*QF*|*RF*)_*x*_ correspond to points along the ESE curve: respectively, the curve maximum (blue), the boundary point nearest the red X, and the red X itself. The gradients of the *UPS*_*D*_ and *UPS*_*I*_ lines are visualised as green and orange slopes, respectively. In example A, one sample in the *FF* exhibits an unexpectedly high expression (yellow) that contradicts the overall negative correlation with the SF. We call this expression value a Divergent observation, corresponding to *D* in Equation (9), and is interpreted as a false positive (FP) according to the ESE flow diagram (Fig S1C). Applying Equation (11) yields an *EP* of 0.049 (Fig S2A, middle panel), indicating that the observed correlation is significant (*EP >* 0) and that the relationship between the QF and RF is unlikely to have arisen by chance.

While this interpretation of the divergent observation as an FP is valid, an alternative hypothesis also exists: the discrepancy may instead reflect false negative (FN) observations in the *FF*. In Fig S2A, these putative FN observations are indicated with red square brackets. We refer to such discrepancies as Overhang (*O*), representing a mismatch in the cardinalities of *RF*_*m*_ and *QF*_*m*_ (or *RF*_*M*_ and *QF*_*M*_). A non-zero *O* implies that no rearrangement of observations can fully eliminate entropy from the conditional relationship. In this case, the minimum observable *O* is 3, and the actual overhang is 4 due to the additional divergent sample.

To quantify the likelihood that overhangs contribute to weakened correlation of a dependent relationship, we define an uncertainty measure analogous to *EP*_*D*_ such that:

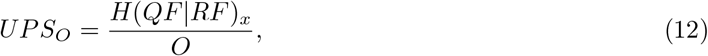

which leads to the overhang error potential;

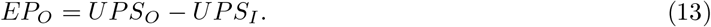

Together, *EP*_*D*_ and *EP*_*O*_ allow us to assess the significance of a correlation and to interpret whether deviations from an ideal relationship are better explained by potential FP or FN observations in the *FF*. Like the ESS, these metrics are symmetric under FF/SF switching. However, as expected they may be asymmetric when comparing *FF* against *SF* observations versus *SF* inverse observations (as illustrated in Fig S2C,D).

By extending the ES framework to incorporate *EP*_*O*_ alongside *EP*_*D*_, we enable more nuanced interpretation of complex relationships in high-dimensional data. This update improves the robustness and interpretability of ES-based tools such as ESFS when applied to complex datasets like scRNA-seq data.

#### Calculating Divergence and Overhang for each Entropy Sort Equation

For any pair of features, we may follow the flow diagram in Fig S1C to identify the reference feature (RF) and query feature (QF) (step **1**.), and to determine which Entropy Sort Equation (ESE) to use for the ES calculations (step **2**.). In step **3**., we compute the observed Divergences (*D*) and Overhangs (*O*) required for ES calculations using a set of defined equations specific to each ESE.

To illustrate how these equations are derived, we take ESE2 as an example. According to Fig S1C,**2**., when the sort direction (*SD* = −1) — meaning non-zero observations are anti-correlated — divergent observations (*D*) occur when non-zero values of the *FF* overlap with non-zero values of the *SF*. Overhang observations (*O*) occur when zero values of the *FF* overlap with zero values of the *SF*. In other words:

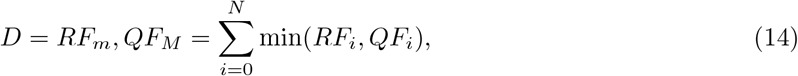

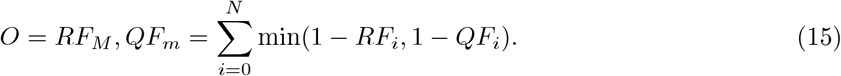

While Equation 15 is valid, directly computing ∑minimum(1 − *RF*_*i*_, 1 − *QF*_*i*_) across all values in a large dataset is computationally expensive, especially because the (1 − *RF*_*i*_) and (1 − *QF*_*i*_) terms hinder efficient use of the sparse structure typical of scRNA-seq data.

To address this, we use a mass balance approach that avoids computing the full element-wise minimum on inverted values. By rearranging Equation 14, we derive an efficient expression for *O*:

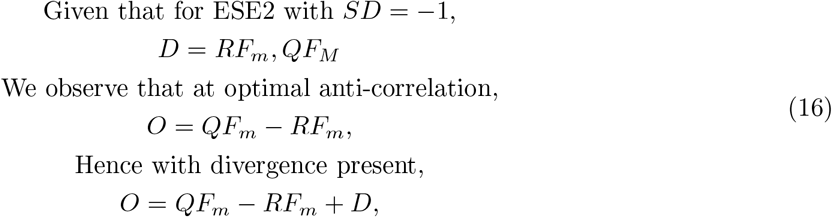

Equation (16) corresponds to the one shown for ESE2 in Fig S1C,**3**. when *SD* = −1.

By applying similar logic to all scenarios in Fig S1C,**3**., we provide a full set of algorithmically efficient equations for calculating *D* and *O* across all possible FF/SF pair types — thereby improving scalability of ES-based analyses on large scRNA-seq datasets.

### Entropy Sorting Feature Selection (ESFS) workflow

In this work, we outline a scRNA-seq analysis workflow termed Entropy Sorting Feature Selection (ESFS). In Fig 1A-C, we provide an overview of the three major Entropy Sorting (ES) derived software that ESFS uses to extract biologically relevant gene expression patterns from scRNA-seq data. Here we provide further detail of the individual steps of these algorithms in relation to the overall ESFS workflow. Reproducible Python Jupyter notebooks for applying ESFS to the datasets used in this manuscript may be found at https://github.com/aradley/ESFS/tree/main/Example_Workflows.

#### Basic quality control

As with most scRNA-seq analysis workflows, it is important to perform basic quality control (QC) to remove poor quality samples and lowly expressed genes. While there can be variation in the approach and parameters applied to different scRNA-seq datasets, the most widely used criteria include relative library size, number of detected genes and fraction of reads mapping to mitochondria-encoded genes or synthetic spike-in RNAs^89^. Having applied a combination of these metrics to remove poor quality cells and genes from the data, we may move to the first stage of the ESFS workflow.

#### Entropy Sorting Gene Set Selection (ES-GSS)

It is widely accepted that subsetting scRNA-seq datasets to include genes enriched for biologically informative expression – and depleted of non-specific or noisy signals – improves downstream analysis. A common approach for this is highly variable gene (HVG) selection. However, due to its univariate nature, HVG selection can lack robustness across methods and may struggle to distinguish between genes that are globally variable and those expressed in distinct cell types^3,5^.

In previous work, we introduced a multivariate alternative, Entropy Sort Feature Weighting (ESFW), which used the ES framework to identify and rank genes that significantly co-regulate across a dataset^5,15^. As part of the ESFS package, we have updated ESFW to incorporate the theoretical improvements to the ES framework outlined in Entropy Sorting theory updates.

ESFW simultaneously computes both a correlation metric (*ESS*) and a correlation significance metric (*EP*) between all gene pairs. For details regarding the calculation of the ESS and EP metrics, see Materials and Methods and Radley et al. 2023, Radley, Boeing, and Smith 2024. Significant pairwise interactions (as determined by *EP* values greater than 0 (Fig S1, Fig S2)) are then used to calculate a weighted node centrality score for each gene (Fig 1i.), such that each gene’s node centrality is give by;

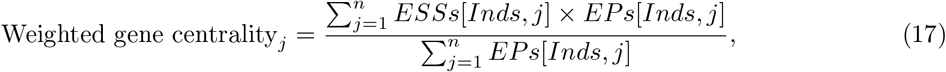

where *j* denotes the column/gene index in the pairwise *ESS* and *EP* matrices, and *Inds* denotes the rows of the *EP* matrix where *EP >* 0 in column *j*. By weighting each value in *ESS* by the corresponding values in *EP*, we mitigate the contribution of uninformative/random correlative patterns to the calculation of gene centralities.

These centrality scores are used to rank genes from most to least connected. The top-ranked genes are then selected for gene set clustering (Fig 1ii.), where the dense *ESS* correlation is converted to a sparse correlation matrix by setting *ESS* values to 0 at all indices where *EP <* 0. This sparse *ESS* correlation matrix is then used to cluster genes into co-expressing sets. This step allows genes with distinct expression patterns to be grouped separately, enriching for biologically informative signal while reducing the impact of technical noise or gene co-expression profiles that are not related to the biological process of interest.

Together, ESFW and subsequent gene set selection make up the Entropy Sorting Gene Set Selection (ES-GSS) algorithm. As we demonstrate in this manuscript, this multivariate approach can improve the resolution of cell states and developmental trajectories in high throughput transcriptomics data.

Selecting the number of top ranked ESFW genes and which set of genes to proceed with for downstream analysis is facilitated by built in ESFS functions that allow the user to easily identify distinct genesets and visualise the resulting UMAP embedding. As with HVG selection in standard Scanpy/Seurat workflows, choosing the number of top-ranked genes is typically an iterative process. To help new users understand how to choose the number of top genes to select, in the Experimental Procedures section we present a sensitivity analyses for the ES-GSS algorithm across each of the datasets used in this manuscript, and on our GitHub repository (https://github.com/aradley/ESFS) we provide workflows for reproducing analysis of each of the 4 datasets in this manuscript.

#### Unbiased cell state marker gene identification

A critical step in scRNA-seq analysis is identifying gene expression signatures that define distinct cell populations. Conventionally, this is achieved by clustering cells using a distance metric, followed by differential expression testing to identify marker genes. However, this approach depends heavily on the chosen clustering resolution, which may either obscure biologically meaningful subpopulations or introduce spurious groupings.

One way to mitigate this issue is to intentionally over-cluster the data and then evaluate all possible combinations of clusters to identify those that best correlate with gene expression profiles. Unfortunately, this combinatorial space is computationally intractable. For example, calculating the Entropy Sorting Score (ESS) for 20,000 genes across all combinations of 100 clusters would require approximately 1.45 *×* 10^28^ CPU years.

Here, we show that the ES framework overcomes this computational bottleneck by reformulating the problem into a tractable sorting task. Specifically, for any pair of features, the ESS can be decomposed into two components: Sort Gain (SG), which captures expression heterogeneity, and Sort Weight (SW), which accounts for magnitude differences. By focusing solely on SG values to rank clusters, and ignoring SWs, we can rapidly evaluate candidate combinations. Using this approach, the previously intractable clustering problem becomes solvable in 100 CPU minutes, rather than 1.45 *×* 10^28^ CPU years.

This performance gain allows us to build a practical algorithm for identifying, for each gene, the combination of clusters in an over-clustered dataset that maximally correlates with its expression profile. We term this algorithm Entropy Sorting Combinatorial Cluster Finder (ES-CCF). These optimised gene-cluster assignments form the basis for Entropy Sorting Find Marker Genes (ES-FMG), which identifies minimal, robust marker genesets from scRNA-seq data.

#### Entropy Sorting Combinatorial Cluster Finder (ES-CCF)

The ability of ES to efficiently determine which cluster combinations maximise a gene’s ESS enables us to over-cluster a dataset—capturing fine-grained local gene expression signatures—without sacrificing the ability to detect broad, global patterns. After first improving scRNA-seq data interpretability using ES-GSS, the ES-CCF algorithm is applied to an intentionally over-clustered dataset (Fig 1iii.).

Over-clustering can be achieved using a variety of approaches. For the mouse colon and early human embryo datasets, we used Leiden clustering from the Scanpy package with a resolution parameter significantly higher than the default value of 1. For the mouse neural tube dataset, where Leiden clustering failed to adequately resolve branching subpopulations within spatial domains, we applied hierarchical clustering with a target of 250 clusters.

Once an over-clustered representation is generated, ES-CCF identifies for each gene the cluster combination that maximises its ESS. To achieve this efficiently, the algorithm computes the SG of each gene with respect to every individual cluster and ranks clusters by descending SG (Fig 1iv.). By focusing on SG alone, this ranking reflects the homogeneity of gene expression within each cluster, independent of cluster size.

This ranking allows us to transform an exponential combinatorial problem—evaluating 2^*n*^ − 1 combinations—into a linear one, requiring only *n* − 1 steps. Stepping through the ranked clusters and computing the cumulative ESS at each step reveals a unimodal curve: the ESS increases until a peak and then decreases (Fig 1v.). The peak ESS corresponds to the optimal cluster combination for that gene (Fig 1vi.), which we refer to as the gene’s max ESS cluster.

After applying ES-CCF to the full gene set, users obtain a max ESS cluster assignment for every gene in the dataset (Fig 1vii.), providing a powerful coarse-grained map of robust gene expression domains across the cellular landscape.

#### Entropy Sorting Find Marker Genes (ES-FMG)

The output of ES-CCF is a two-dimensional matrix mapping each gene to its corresponding max ESS cluster across all samples in the dataset (Fig 1vii.). Each max ESS cluster provides a coarse-grained approximation of the region in which a gene’s expression is most strongly enriched. Our goal is to identify a small set of these gene-cluster pairs that together capture distinct, non-overlapping gene expression profiles across the combinatorial cluster space. Because each max ESS cluster is already linked to a gene, this process simultaneously yields an optimised, non-redundant marker gene set.

To do this, we first compute the pairwise ESS between all max ESS clusters and all genes in the dataset, resulting in a square matrix (max ESS cluster × gene), which quantifies the correlation between each coarse-grained cluster and gene expression profiles (Fig 1viii.).

From this matrix, we aim to select a set of *N* clusters (and their corresponding genes) that together capture maximally distinct expression patterns. Specifically, we seek clusters where each selected gene has strong correlation with its own cluster and minimal correlation with the others. This is formalised by the following objective function:

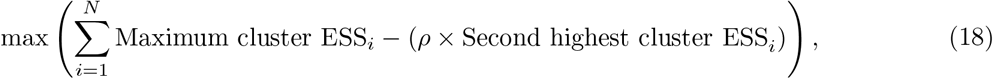

Here, *ρ* is a tunable parameter that controls the penalty for redundancy. A *ρ* of 1 heavily penalises overlap in expression patterns between marker genes, encouraging maximal distinctness. A *ρ* of 0 imposes no penalty, allowing selection of genes with overlapping expression profiles. Intermediate values enable flexible control over the trade-off between distinctiveness and redundancy, which is especially useful for capturing gradient-like or overlapping cellular states. Sensitivity analyses for *ρ* are provided in the Experimental Procedures section.

To optimise this objective, the following iterative procedure is applied:

1. Initialise a set of *n* clusters by random selection.
2. Subset the full correlation matrix to an *n × n* submatrix corresponding to the selected clusters and their associated genes (Fig 1x.).
3. For each cluster in the set, evaluate all possible replacements with unused clusters, calculating the new value of (18).
4. Replace the cluster that yields the greatest improvement in the objective function.
5. Repeat steps 2–4 until no further improvement is observed.

A visualisation of an optimised *n* = 20 subset is shown in Fig 1x., with green highlights denoting the maximum ESS values along the diagonal, and red boxes marking the second-highest ESS values for each potential marker gene, which are penalised in the optimisation.

The result of this process is a minimal set of *n* clusters and corresponding marker genes that together span the dominant, non-redundant expression programs present in the combinatorial cluster space. Visualising these optimised marker genes reveals distinct cellular populations with robust expression profiles (Fig 1xi.). As *n* is typically much smaller than the full gene set, this procedure greatly simplifies the identification of biologically relevant marker genes in complex and heterogeneous scRNA-seq datasets (Fig 1xii.).

