## Supplemental Material for "Entropy Sorting Feature Selection: information-theoretic gene set identification improves single-cell RNA sequencing data interpretability"

### SUPPLEMENTARY MATERIAL

| Col1 | Col2 | Col2 | Col3 |
|---|---|---|---|
| 1 | 6 | 87837 | 787 |
| 2 | 7 | 78 | 5415 |
| 3 | 545 | 778 | 7507 |
| 4 | 545 | 18744 | 7560 |
| 5 | 88 | 788 | 6344 |

Table S1: **A placeholder table for the mouse neural tube cell type ranked gene list table which has been uploaded with this submission.**

| Nomenclature | Description |
|---|---|
| ES | Entropy Sorting - A mathematical framework originally derived in Radley et al. 2023. |
| ESFS | Entropy Sorting Feature Selection - A modular software developed for high dimensional transcriptomics data analysis. |
| $N$ | The number of observations/samples comprising each feature. For scRNA-seq data this is equal to the number of cells in the counts matrix. |
| Feature | An individual, measurable property of an observed phenomenon. In scRNA-seq data, each gene is a feature measured across $N$ samples (cells), where each sample records the number of observed transcripts. For ES, feature activity in each sample is normalised to the range [0,1], with 0 indicating absence and 1 indicating maximal activity. Inactivity is defined inversely, where 0 represents maximal inactivity and 1 represents no inactivity. |
| $_m$ | Minority states - $_m$ denotes a scalar value of the sum of the less common observation type between observed activities and inactivities in a given feature. We determine the minority states for a given feature via equations (1) and (2) in the Materials and Methods. |
| $_M$ | Majority states - $_M$ denotes a scalar value of the sum of the more common observation type between observed activities and inactivities in a given feature. We determine the majority states for a given feature via equations (1) and (2) in the Materials and Methods. |
| RF and QF | For any pair of features (i.e. genes), to utilise the ES framework, one feature must be designated the Reference Feature (RF) and the other the Query Feature (QF). According to the maximum entropy principal, the QF is always the feature with the larger number of observer minority states ($QF_m > RF_m$). |
| $RF_m$ and $QF_m$ | The sum of minority states ($_m$) in the RF or QF. It follows that $RF_m + RF_M = N$. |
| $RF_M$ and $QF_M$ | The sum of majority states ($_M$) in the RF or QF. It follows that $QF_m + QF_M = N$. |
| $QF_m, RF_m$, $QF_M, RF_m$, $QF_m, RF_M$, $QF_M, RF_M$ | Scalar values indicating the total co-occurrence/overlap of minority states ($_m$) and/or majority states ($_M$) between the $N$ samples of the QF and RF. Algorithmically for any feature pair each of these variable is given by, $\sum_{i=0}^{N} \min(RF_i, QF_i)$, and it is the composition of the FF and SF that determines whether $\sum_{i=0}^{N} \min(RF_i, QF_i)$ equals $QF_m, RF_m$, $QF_M, RF_m$, $QF_m, RF_M$ or $QF_M, RF_M$. |
| $H(QF\|RF)$ | The conditional entropy of the QF conditioned on the RF. |
| ESE | Entropy Sort Equation - There are 4 Entropy Sort Equations (Fig S1B, ESE 1-4), each presenting a re-arrangment of conventional probabilistic conditional entropy into a sorting problem. Fig S1C presents a flow diagram for determining which ESE should be applied for any given pair of features. |
| $ESS$ | Entropy Sort Score - A metric derived from the ES framework to quantify the correlation between two features. |
| $SW$, $SG$ and $SD$ | Sort Weight, Sort Gain and Sort Direction - When multiplied together, these three quantities derived from the ESE equal the ESS - $ESS = SW \times SG \times SD$. |
| $EP$ | Error Potential - A metric derived from the ES framework to determine whether the correlation between two features is significant. |
| $FF$ and $SF$ | Algorithmically, when calculating ES metrics it is useful to focus on one particular feature, the Fixed Feature ($FF$) and compare it pairwise against every other Secondary Feature ($SF$). See Fig S1C for algorithmic demonstration. |
| $D$ and $O$ | Divergence and Overhang - For any pair of features, Divergence and Overhang capture potential false negative or false positive values that weaken the hypothesised dependence between the features. |
| $UPS_I$ | Uncertainty Per Sample when a pair of features are maximally independent from one another. |
| $UPS_D$ and $UPS_O$ | Uncertainty Per Sample due to the presence of Divergent ($D$) or Overhang ($O$) values that weaken the dependent relationship between a pair of feature. |
| $EP_D$ and $EP_O$ | The Error Potential between a pair of features due to the presence of Divergent ($D$) or Overhang ($O$) observations. |

Table S2: **Table of Entropy Sorting nomenclature.**

**A.**

**Conventional conditional shannon entropy equation**

$$H(QF|RF) = \frac{RF_m}{RF_m + RF_M}\left(-\frac{QF_m,RF_m}{RF_m}\log\left(\frac{QF_m,RF_m}{RF_m}\right) - \frac{QF_M,RF_m}{RF_m}\log\left(\frac{QF_M,RF_m}{RF_m}\right)\right) + \frac{RF_M}{RF_m + RF_M}\left(-\frac{QF_m,RF_M}{RF_M}\log\left(\frac{QF_m,RF_M}{RF_M}\right) - \frac{QF_M,RF_M}{RF_M}\log\left(\frac{QF_M,RF_M}{RF_M}\right)\right)$$

**B.**

**ESE 1: x = $QF_m,RF_m$**

$$H(QF|RF) = \frac{RF_m}{RF_m + RF_M}\left(-\frac{QF_m,RF_m}{RF_m}\log\left(\frac{QF_m,RF_m}{RF_m}\right) - \frac{RF_m - QF_m,RF_m}{RF_m}\log\left(\frac{RF_m - QF_m,RF_m}{RF_m}\right)\right) + \frac{RF_M}{RF_m + RF_M}\left(-\frac{QF_m - QF_m,RF_m}{RF_M}\log\left(\frac{QF_m - QF_m,RF_m}{RF_M}\right) - \frac{RF_M - QF_m + QF_m,RF_m}{RF_M}\log\left(\frac{RF_M - QF_m + QF_m,RF_m}{RF_M}\right)\right)$$

**ESE 2: x = $QF_M,RF_m$**

$$H(QF|RF) = \frac{RF_m}{RF_m + RF_M}\left(-\frac{RF_m - QF_M,RF_m}{RF_m}\log\left(\frac{RF_m - QF_M,RF_m}{RF_m}\right) - \frac{QF_M,RF_m}{RF_m}\log\left(\frac{QF_M,RF_m}{RF_m}\right)\right) + \frac{RF_M}{RF_m + RF_M}\left(-\frac{RF_M - QF_M + QF_M,RF_m}{RF_M}\log\left(\frac{RF_M - QF_M + QF_M,RF_m}{RF_M}\right) - \frac{QF_M - QF_M,RF_m}{RF_M}\log\left(\frac{QF_M - QF_M,RF_m}{RF_M}\right)\right)$$

**ESE 3: x = $QF_m,RF_M$**

$$H(QF|RF) = \frac{RF_m}{RF_m + RF_M}\left(-\frac{QF_m - QF_m,RF_M}{RF_m}\log\left(\frac{QF_m - QF_m,RF_M}{RF_m}\right) - \frac{RF_m - QF_M + QF_m,RF_M}{RF_m}\log\left(\frac{RF_m - QF_M + QF_m,RF_M}{RF_m}\right)\right) + \frac{RF_M}{RF_m + RF_M}\left(-\frac{QF_m,RF_M}{RF_M}\log\left(\frac{QF_m,RF_M}{RF_M}\right) - \frac{RF_M - QF_m,RF_M}{RF_M}\log\left(\frac{RF_M - QF_m,RF_M}{RF_M}\right)\right)$$

**ESE 4: x = $QF_M,RF_M$**

$$H(QF|RF) = \frac{RF_m}{RF_m + RF_M}\left(-\frac{RF_m - QF_M + QF_M,RF_M}{RF_m}\log\left(\frac{RF_m - QF_M + QF_M,RF_M}{RF_m}\right) - \frac{QF_M - QF_M,RF_M}{RF_m}\log\left(\frac{QF_M - QF_M,RF_M}{RF_m}\right)\right) + \frac{RF_M}{RF_m + RF_M}\left(-\frac{RF_M - QF_M,RF_M}{RF_M}\log\left(\frac{RF_M - QF_M,RF_M}{RF_M}\right) - \frac{QF_M,RF_M}{RF_M}\log\left(\frac{QF_M,RF_M}{RF_M}\right)\right)$$

**C.**

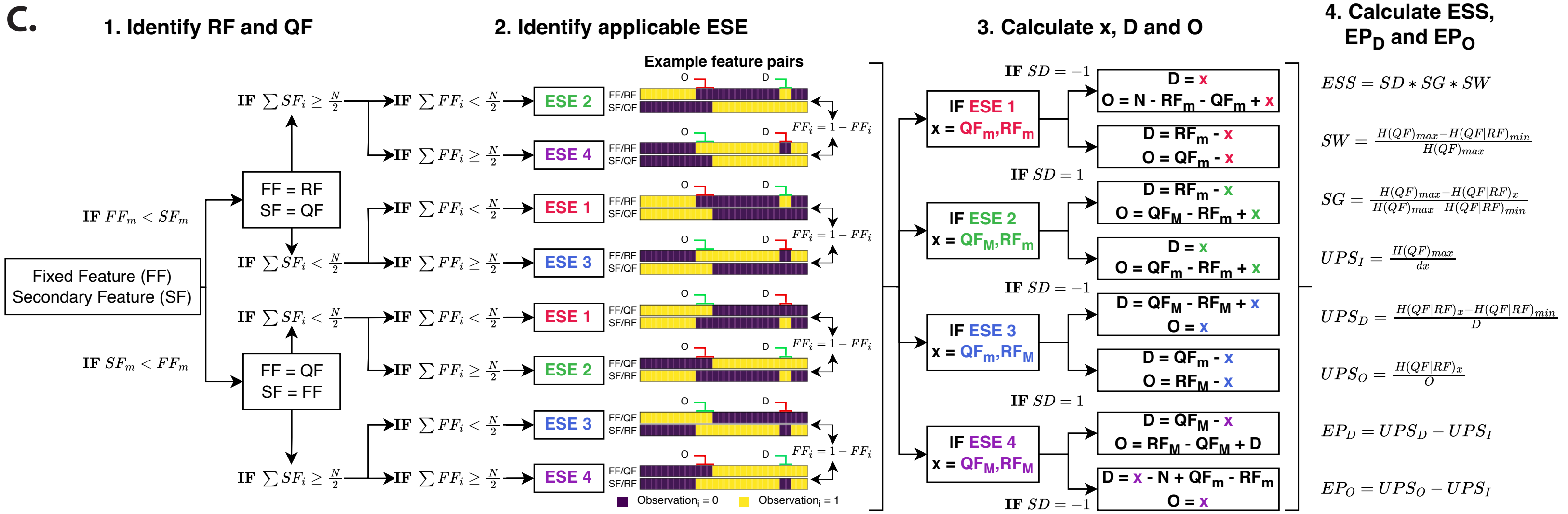


Figure S1: **Updating the ES framework to use all four ESE arrangements. A.** The conventional probabilistic conditional entropy (CE) formulation with entropy sorting notation. $RF_m$ - total reference feature minority states, $RF_M$ - total reference feature majority states, $QF_m$ - total query feature minority states, $QF_M$ - total query feature majority states, $QF_m, RF_m$ - total co-occurrence of RF and QF minority states, $QF_M, RF_m$ - total co-occurrence of RF minority and QF majority states, $QF_m, RF_M$ - total co-occurrence of RF majority and QF minority states, $QF_M, RF_M$ - total co-occurrence of RF and QF majority states. **B.** By substitution of one of the 4 joint variables into the conventional CE equation we can create 4 new Entropy Sort Equations (ESEs) where only one variable remains. **C.** Flow diagram outlining how logically determine which ESE calculations are suitable for any given pair of random variables. Although Entropy Sort Scores ($ESSs$) and Error Potentials ($EPs$) are symmetrical, algorithmically is it practical to designate one feature, the Fixed Feature (FF), as being compared against a Secondary Feature (SF). 1. Identify whether the FF is the RF or QF according to minority state cardinalities. 2. Identify which of $ESE$ is suitable for the observed FF/SF pair. 3. Based on which $ESE$ is applicable, use the correct equations to identify the feature overlap ($x$) and calculate the Divergence ($D$) and Overhang ($O$). 4. Having obtained values for all the required variables, calculate the $ESS$, Divergent Error Potential ($EP_D$) and Overhang Error Potential ($EP_O$).

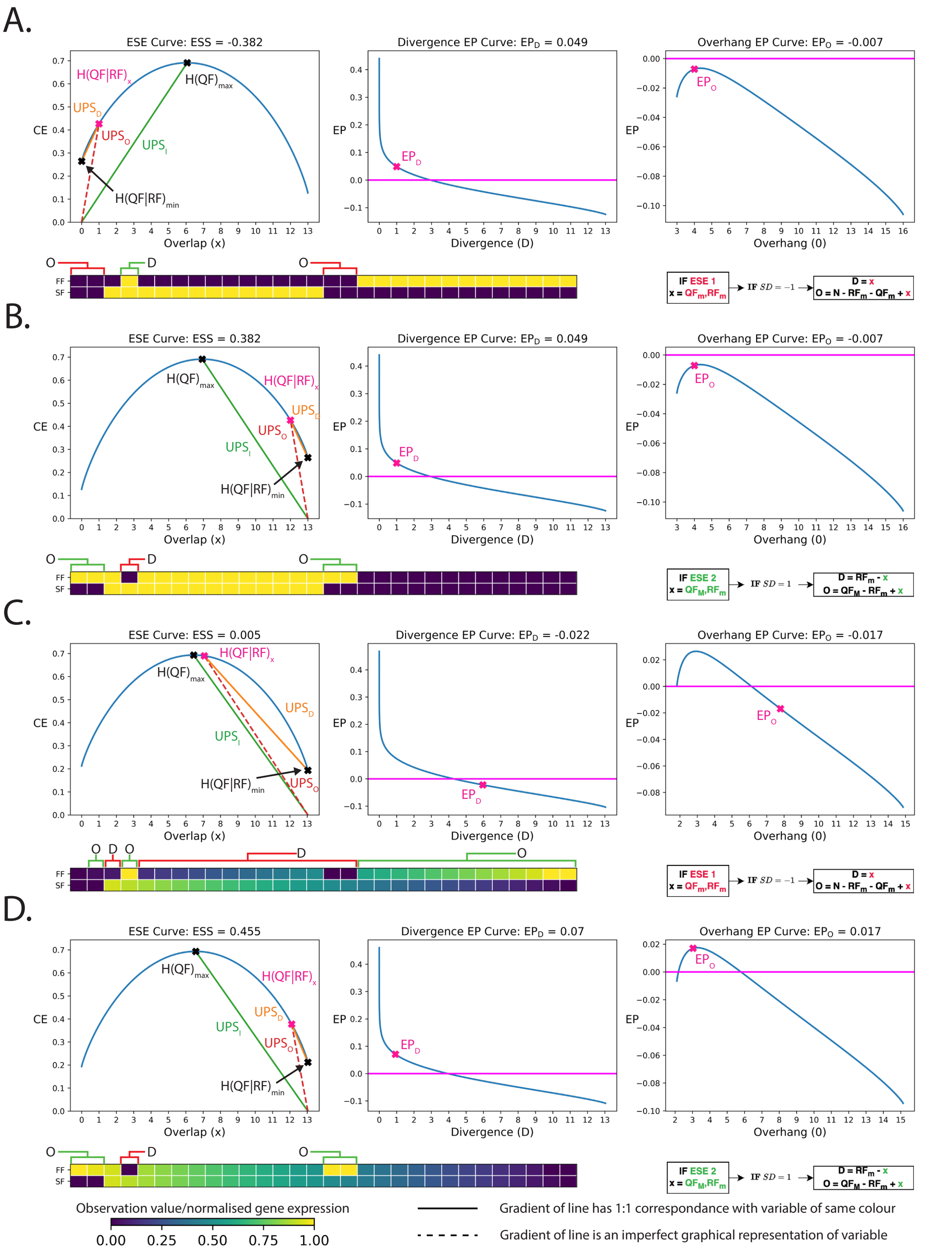


Figure S2: **Inverse Entropy Sorting calculations ensure that random variable correlations and anti-correlations are correctly identified.** Each panel shows an example Fixed Feature (FF)/ Query Feature (QF) pair of random variables (blue and yellow samples). Square brackets highlight observed Divergent (D) and Overhang (O) samples. Green square brackets indicate the sample in the FF is suggested as a false positive value and red indicates a false negative value. On the ESE curves, $H(QF|RF)$ = observed conditional entropy between the QF and RF, $H(QF)_max$ = the maximum independent entropy of the QF, $H(QF|RF)_min$ = the optimal minimum conditional entropy between the QF and RF, $UPS_D$ = the observed uncertainty per divergent sample, $UPS_O$ = the observed uncertainty per overhang sample and $UPS_I$ = the uncertainty per sample at feature independence. For the Error Potential plots, $EP_D$ = the EP of the divergent samples and $EP_O$ = the EP of the overhang samples. The horizontal purple line indicates EP $= 0$. **A,B.** Example showing that for discretised data, the $ESS$ and $EP$ values are not effected by inverting the values of the FF. **C,D.** Example showing that for continuous data, there are cases where in order to robustly identify correlation or anti-correlation between random the FF and SF, we must calculation for the observed states and the inverse FF states.

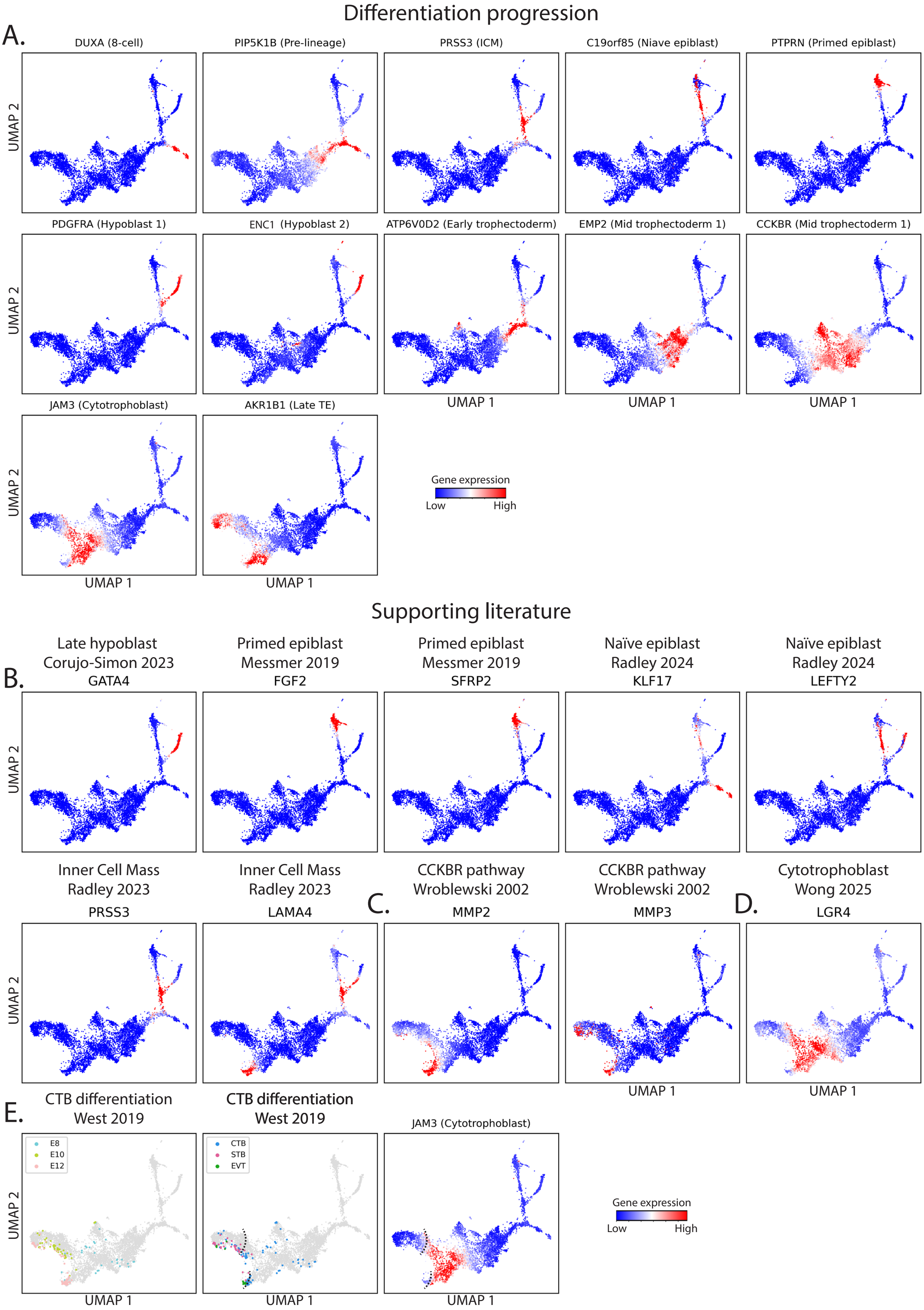


Figure S3: **ESFS reveals distinct gene expression profiles during early human embryogenesis. A.** Individual expression plots for genes highlighted in Fig 2I. **B-D.** Genes expression profiles of genes found in the scientific literature to support the expression dynamics identified by our unsupervised ESFS workflow. **E.** Projecting annotated scRNA-seq data from West et al. 2019's analysis of peri-implantation TE differentiation supports ES-FMG identifying *JAM3* and a CTB marker gene. Black dashed lines indicate a boundary between CTB samples and STB/EVT samples that coincides with the downregulation of *JAM3*.

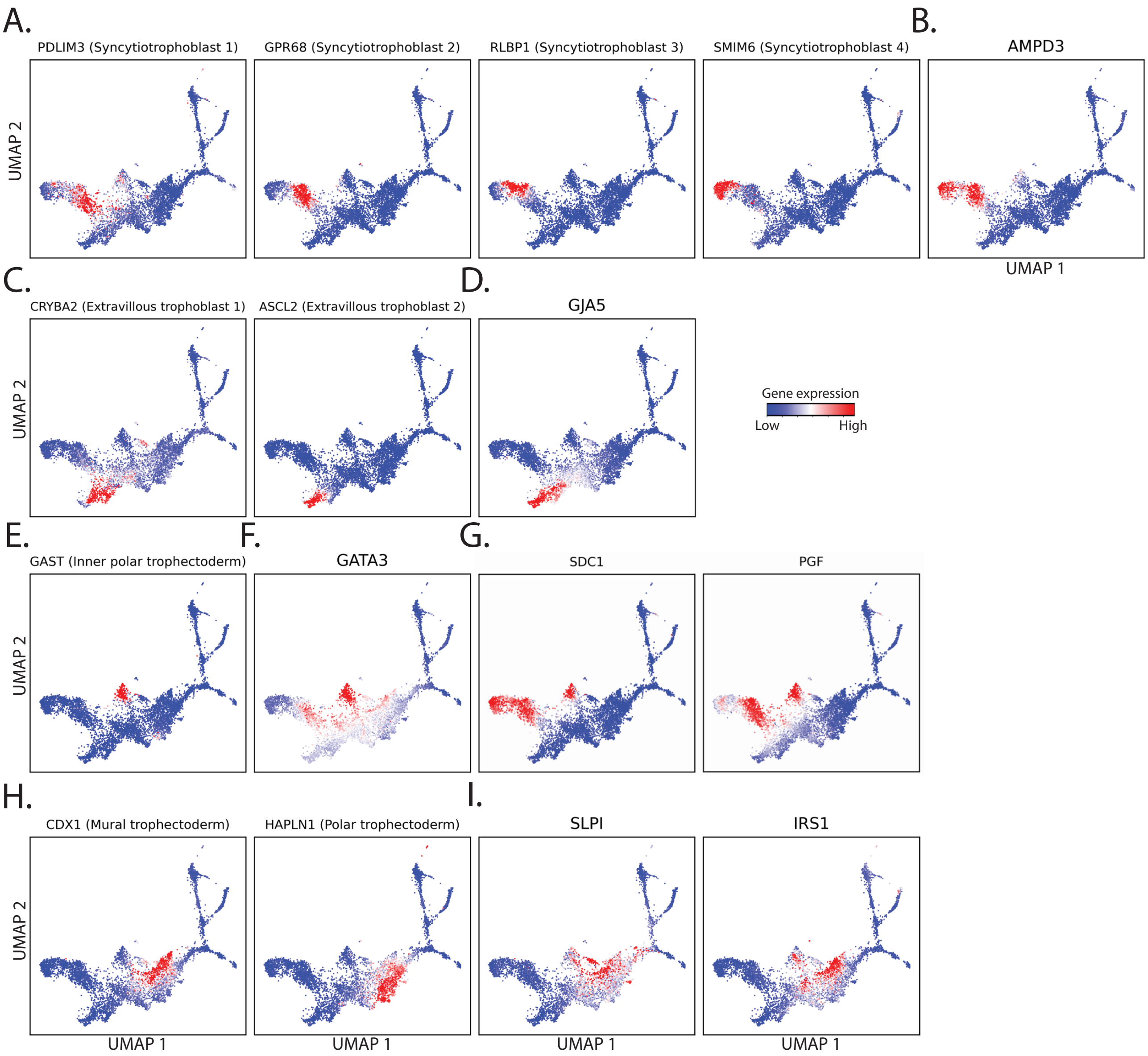


Figure S4: **Individual gene expression profiles of genes characterising trophectoderm diversity in peri-implantation human embryos. A,B.** Marker genes identified by ES-FMG as capturing sub-populations of syncytiotrophoblast (STB) cells. **C,D.** Marker genes identified by ES-FMG as capturing sub-populations of extravillous trophoblast (EVT) cells. **E.** Marker genesidentified by ES-FMG as capturing an inner polar trophectoderm population suggested by Corujo-Simon et al. 2024 as being $GATA3^+$ (F.) in E6-7 human embryos. **G.** Genes validated as polar trophectoderm markers by Liu et al. 2022 via immunofluorescence staining. **H.** Genes identified by ES-FMG as E6-7 mural and polar trophectoderm marker genes. **I.** Genes identified by ES-FMG as being expressed in both mural trophectoderm and the proposed $GAST^+$ inner polar trophectoderm population.

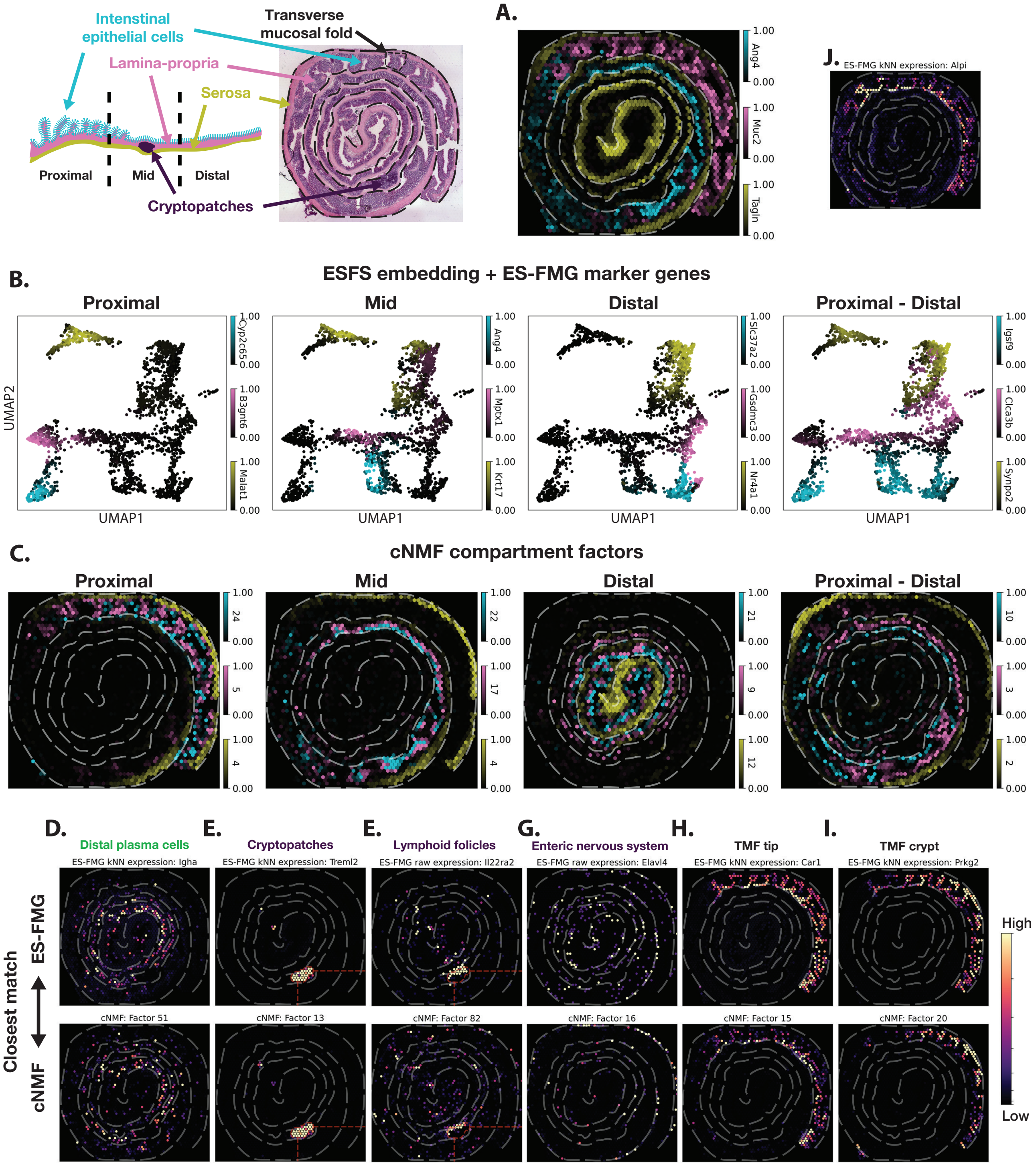


Figure S5: **Cell type diversity in the mouse colon. A.** Spatial visualisation of known cell type marker genes show that they are also spatially restricted. **B.** ES-FMG cell state marker genes are clearly stratified in the UMAP space. **C.** Most similar cNMF factors to cell type/compartment genes identified by ES-FMG in Fig 3F. Colour bars match cell type colours in panel A. **D-G.** Additional cell states identified by Parigi et al. 2022 are also found by ES-FMG and cNMF. **H-J.** Additional cell states identified by ES-FMG.

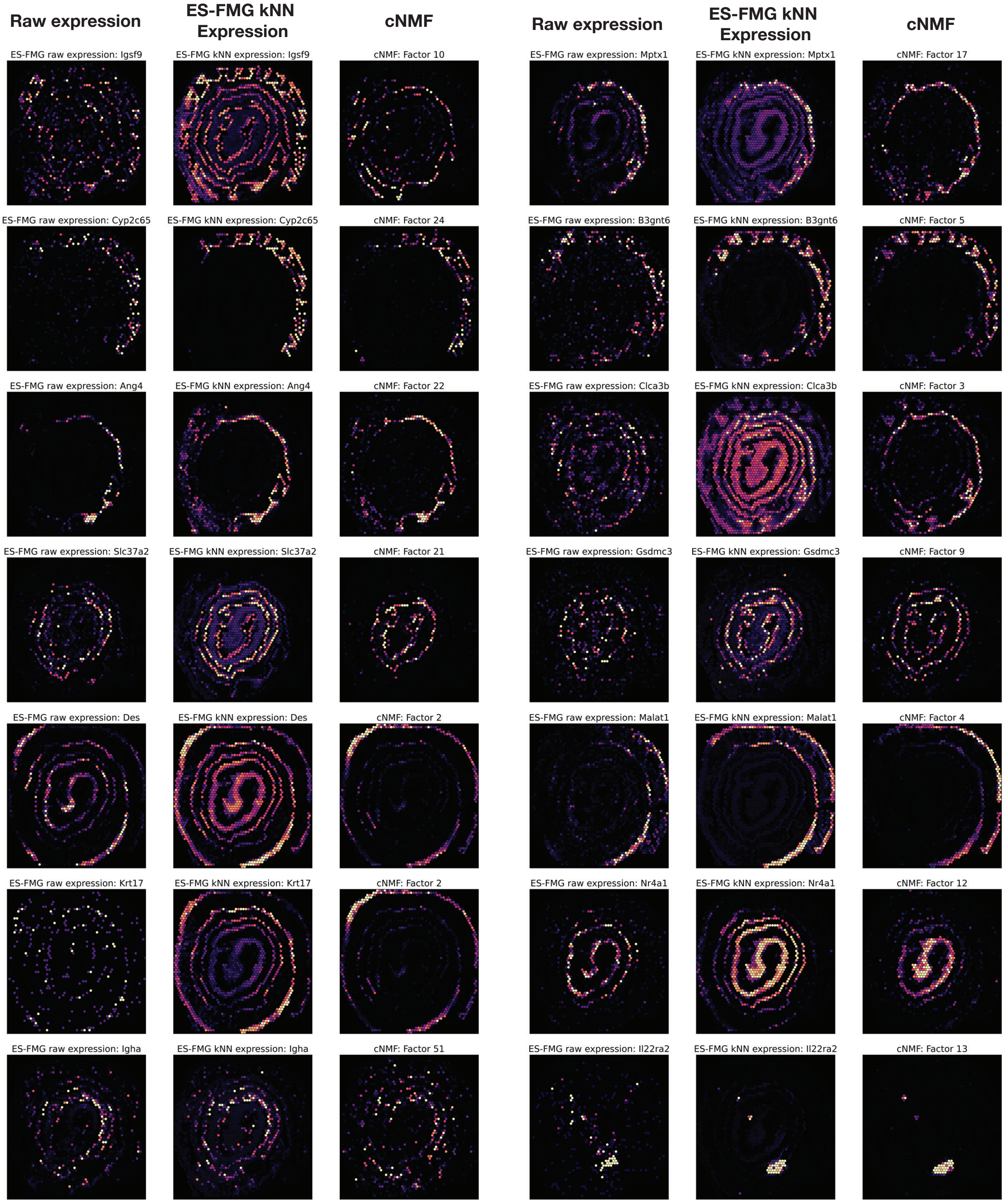


Figure S6: **Spatial transcriptomics plots of the raw gene expression, ES-FMG k-nearest neighbour (kNN) smoothed expression and most similar cNMF factors for genes discussed in main text.**

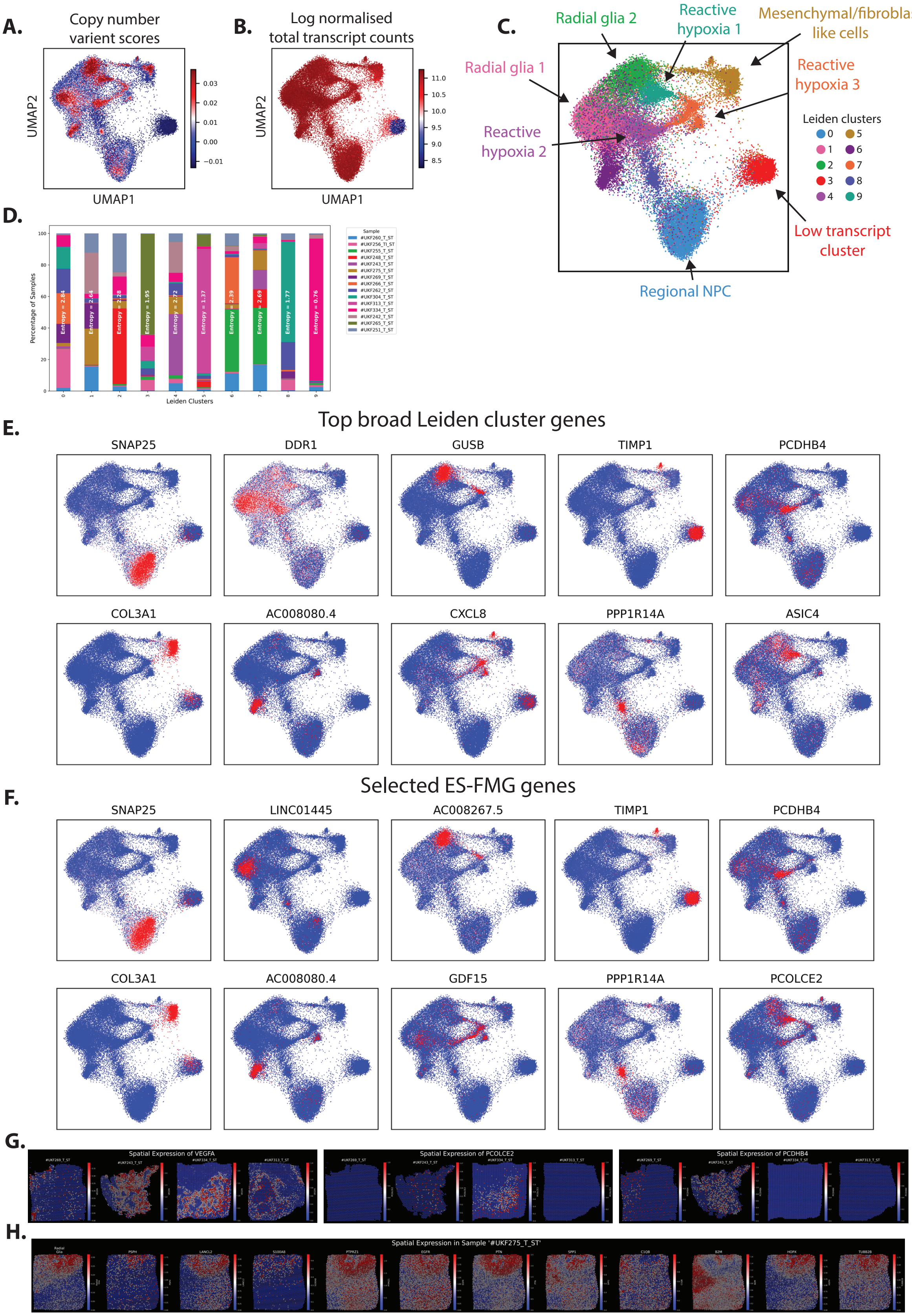


Figure S7: **ESFS captures broad expression programs of glioblastomas. A.** Copy number variant scores show malignant spots are concentrated in the centre of dense spot populations. **B.** Total transcript counts of individual spots reveals a cluster of low expressing/low quality spots. **C.** Default Leiden clustering (Resolution = 1) identifies broad clusters of spots. **D.** Tumour sample heterogeneity within each broad Leiden cluster varies significantly, distinct expression states are poorly conserved across tumours. **E, F.** Top ranked genes from broad Leiden clusters (E.) are closely matched by genes unbiasedly identified via ES-FMG (F.). **G.** Marker genes for distinct Leiden clusters may show spatial restriction in some tumours but be heterogeneously or non-expressed in others. **H.** Activity scores for the radial glia program identified by Ravi et al. 2022 (left most plot) obscure meaningful expression heterogeneity of genes that contribute to the radial glia gene set (11 example genes shown).

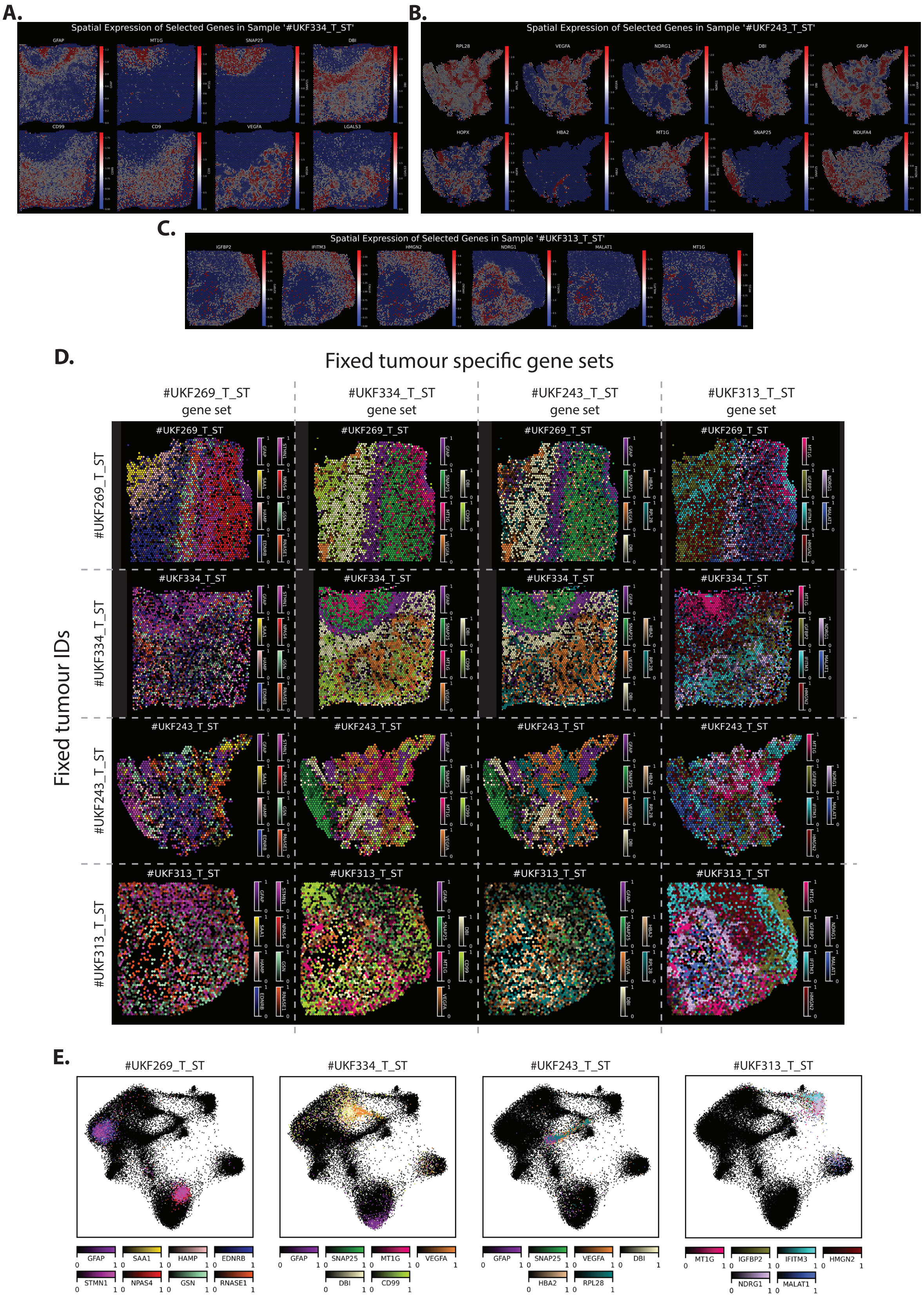


Figure S8: **ESFS streamlines the identification of micro-environments in glioblastomas. A-C.** Selected genes from the 200 ES-FMG genes that show spatially restricted expression in tumours #UKF334_T_ST (A.), #UKF243_T_ST (B.) and #UKF313_T_ST (C.). **D.** Taking gene sets that reveal tumour micro-environments in a particular tumour (diagonal) and visualising their expression in other tumours (off-diagonal) demonstrates the heterogenous nature of glioblastomas and the need for unbiased software that aid marker gene identification. **E.** Visualising tumour specific gene expression profiles in the UMAP space re-affirms that identification of tumour specific expression profiles benefits from ES-CCF and ES-FMGs ability to efficiently search a combinatorial cluster space.

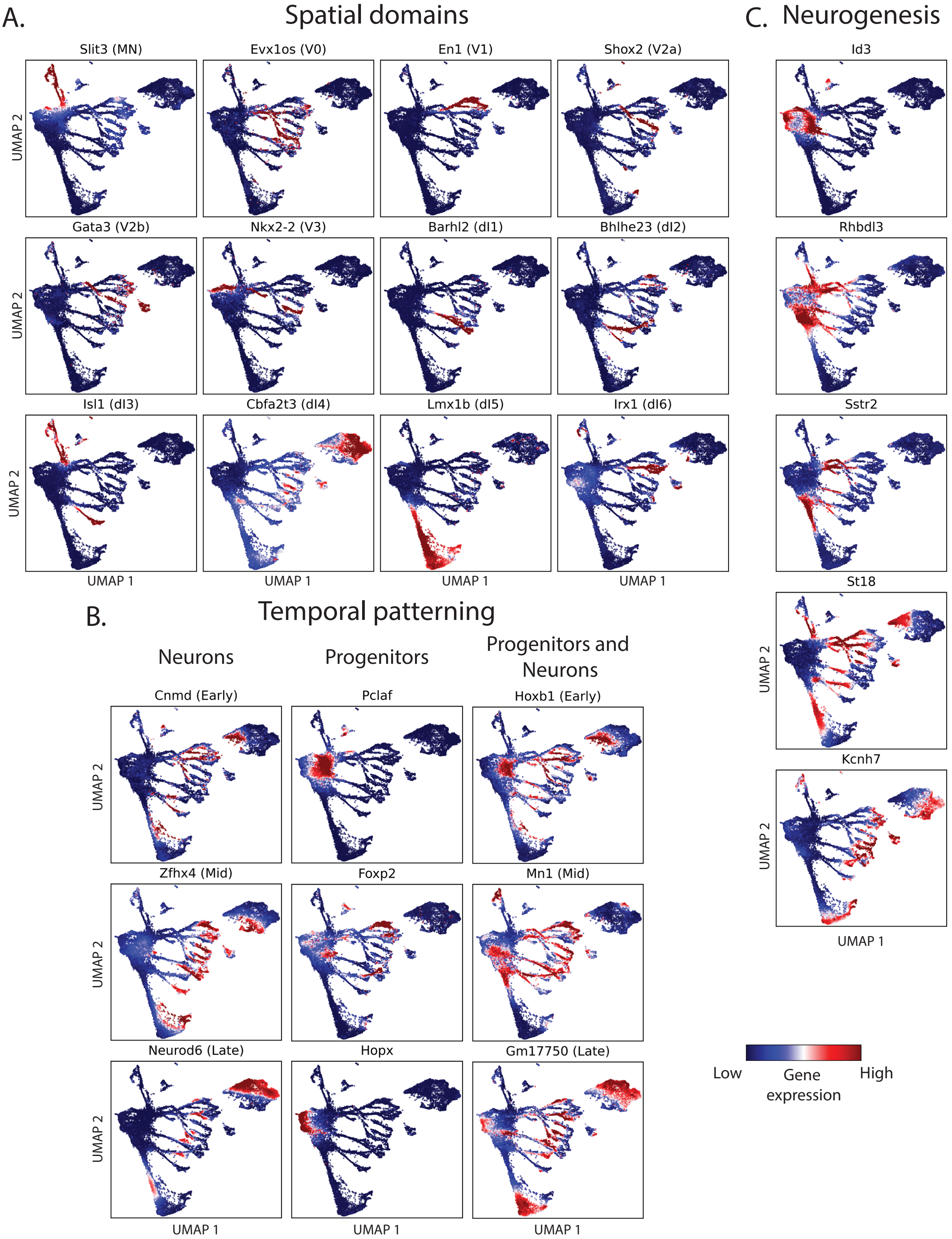


Figure S9: **Individual gene expression profiles of neural tube spatial, temporal and neurogenesis marker genes identified ES-FMG.**

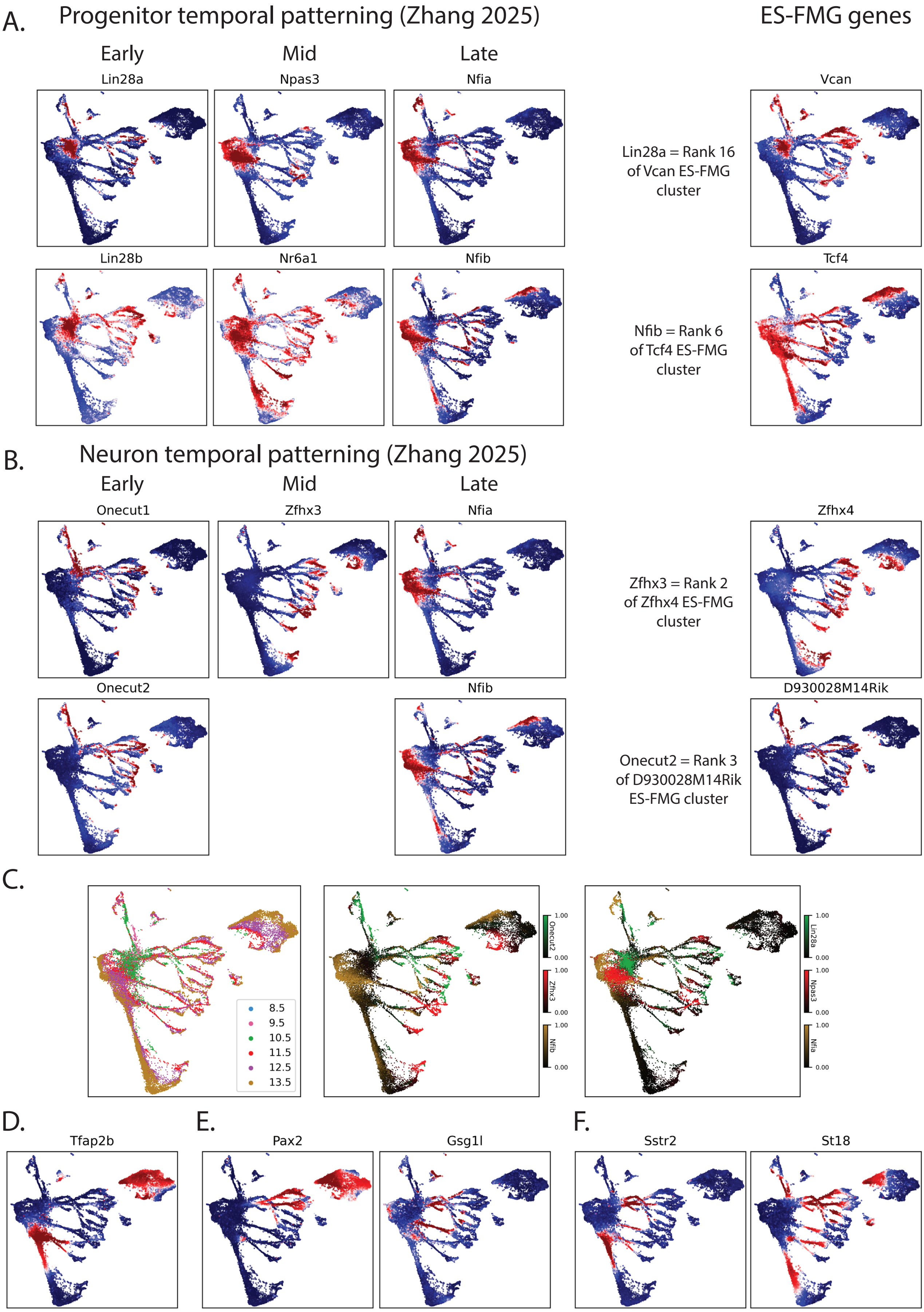


Figure S10: **Marker genes identified from the literature as relating to NT temporal patterning (A-C.) and NT intermediate neurogenesis populations (D-F.)** The ES-FMG gene column that genes identified in the literature have high ranked similarities to genes identified by the ES-FMG algorithm.

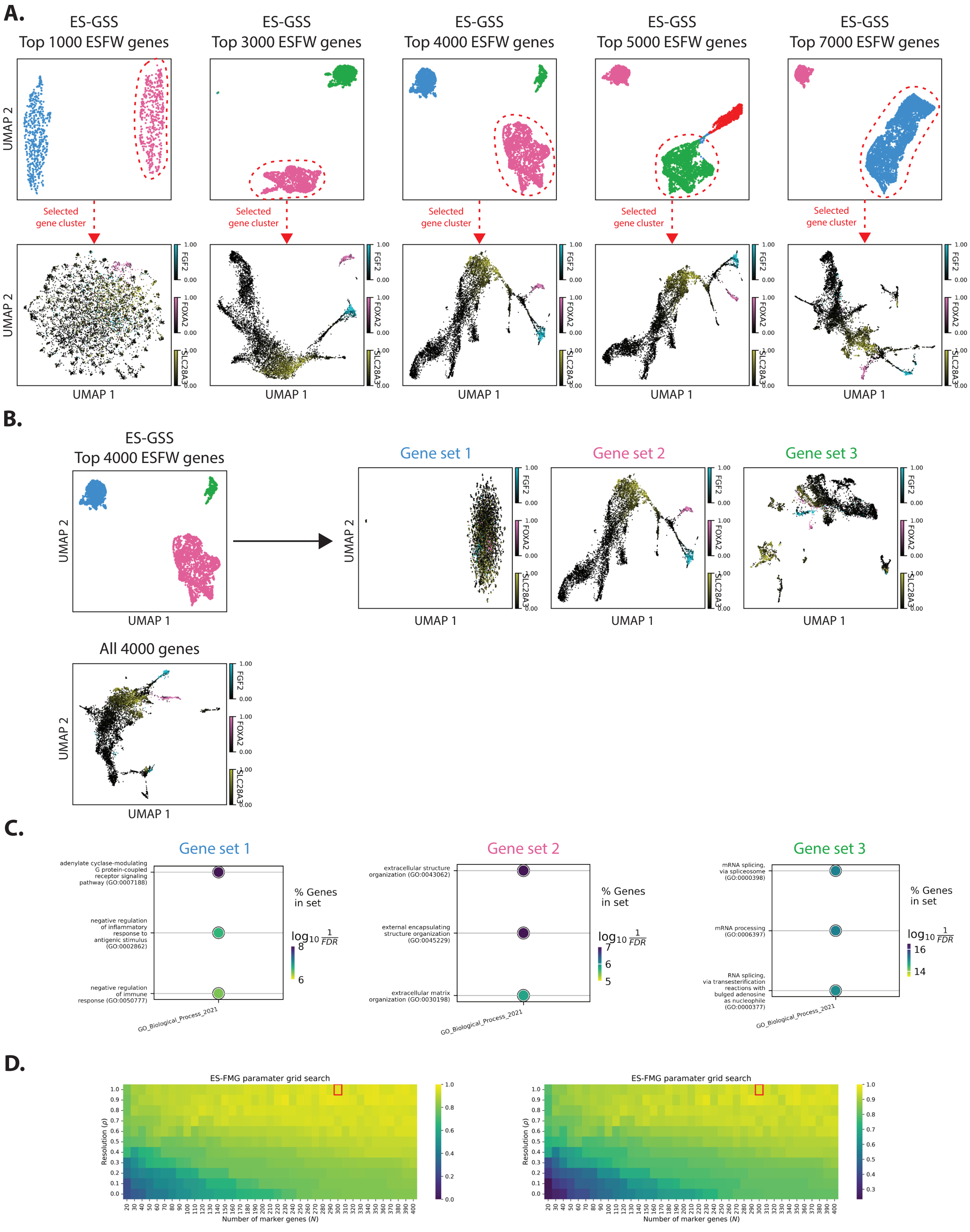


Figure S11: **ESFS parameter selection sensitivity analysis - Early human embryo scRNA-seq. A.** Visualisation of the effect of changing the number of top ranked genes used during gene clustering on the resulting cell UMAP embedding. Gene rankings are determined via entropy sorting feature weighting. **B.** Visualisation of gene cluster choice on the resulting cell UMAP embedding for the number of top ranked genes used in the main manuscript. **C.** Top gene ontology terms obtained via gene set enrichment analysis of biological processes for each of the gene clusters presented in B.. **D.** Sensitivity analysis of parameter selection for the ES-FMG algorithm. A grid search of the number of marker genes to optimise for $(N)$ and the resolution parameter $(\rho)$ shows the expression profiles captured quickly plateau to values close to 1. Red box indicates parameters used in the main manuscript.

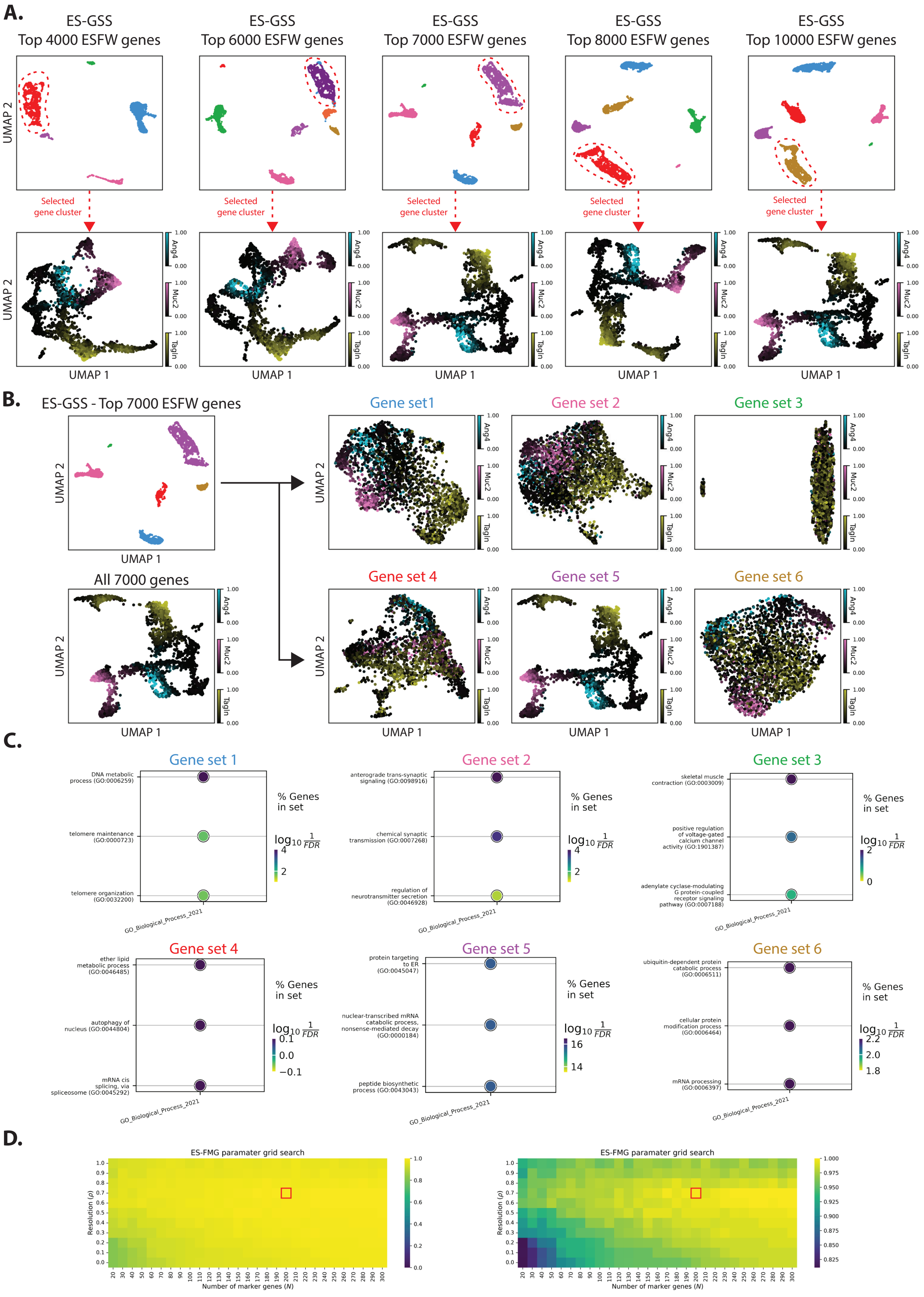


Figure S12: **ESFS parameter selection sensitivity analysis - Mouse colon spatial transcriptomics. A.** Visualisation of the effect of changing the number of top ranked genes used during gene clustering on the resulting cell UMAP embedding. Gene rankings are determined via entropy sorting feature weighting. **B.** Visualisation of gene cluster choice on the resulting cell UMAP embedding for the number of top ranked genes used in the main manuscript. **C.** Top gene ontology terms obtained via gene set enrichment analysis of biological processes for each of the gene clusters presented in B.. **D.** Sensitivity analysis of parameter selection for the ES-FMG algorithm. A grid search of the number of marker genes to optimise for ($N$) and the resolution parameter ($\rho$) shows the expression profiles captured quickly plateau to values close to 1. Red box indicates parameters used in the main manuscript.

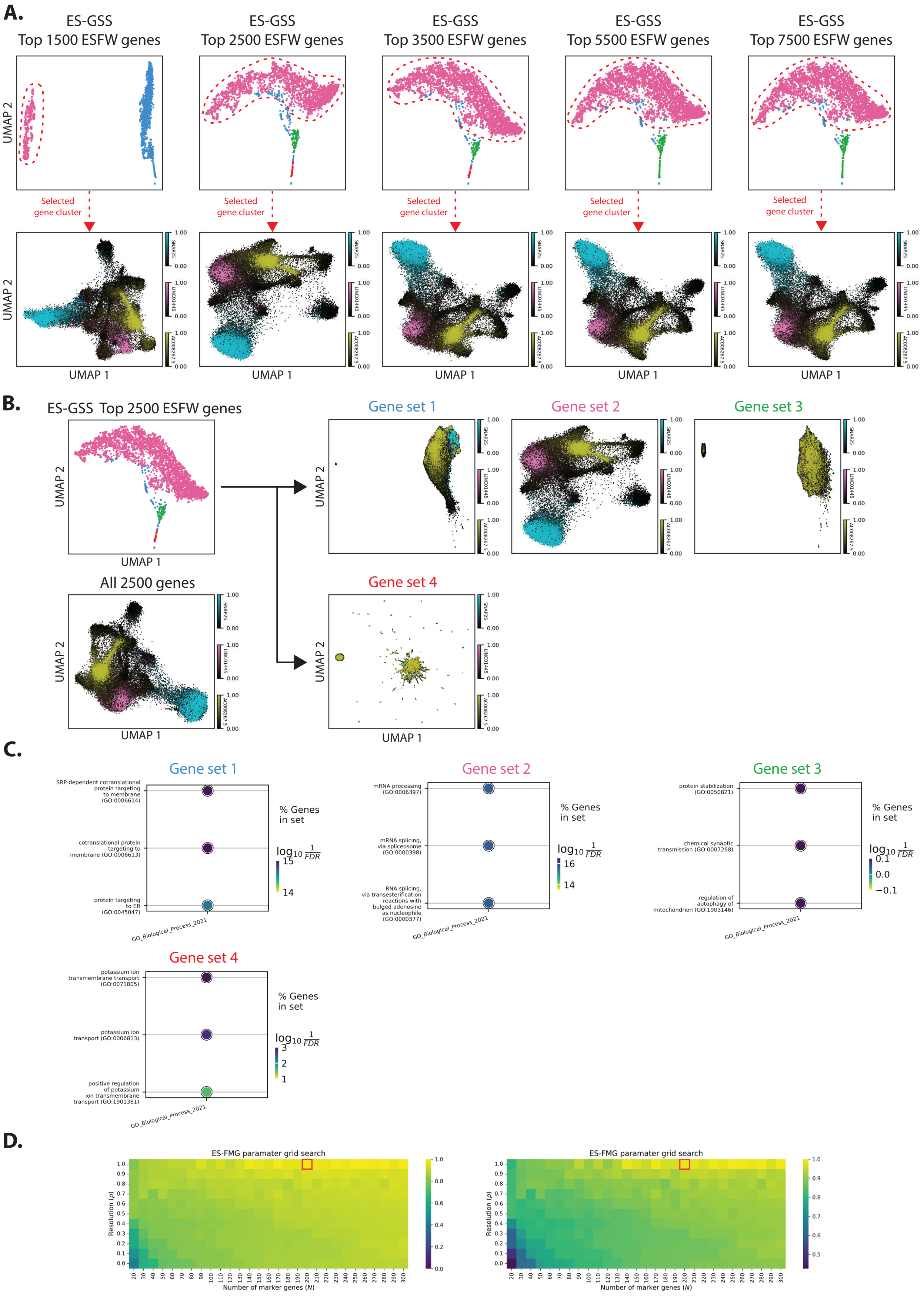


Figure S13: **ESFS parameter selection sensitivity analysis - Human glioblastoma spatial transcriptomics.** **A.** Visualisation of the effect of changing the number of top ranked genes used during gene clustering on the resulting cell UMAP embedding. Gene rankings are determined via entropy sorting feature weighting. **B.** Visualisation of gene cluster choice on the resulting cell UMAP embedding for the number of top ranked genes used in the main manuscript. **C.** Top gene ontology terms obtained via gene set enrichment analysis of biological processes for each of the gene clusters presented in B.. **D.** Sensitivity analysis of parameter selection for the ES-FMG algorithm. A grid search of the number of marker genes to optimise for $(N)$ and the resolution parameter $(\rho)$ shows the expression profiles captured quickly plateau to values close to 1. Red box indicates parameters used in the main manuscript.

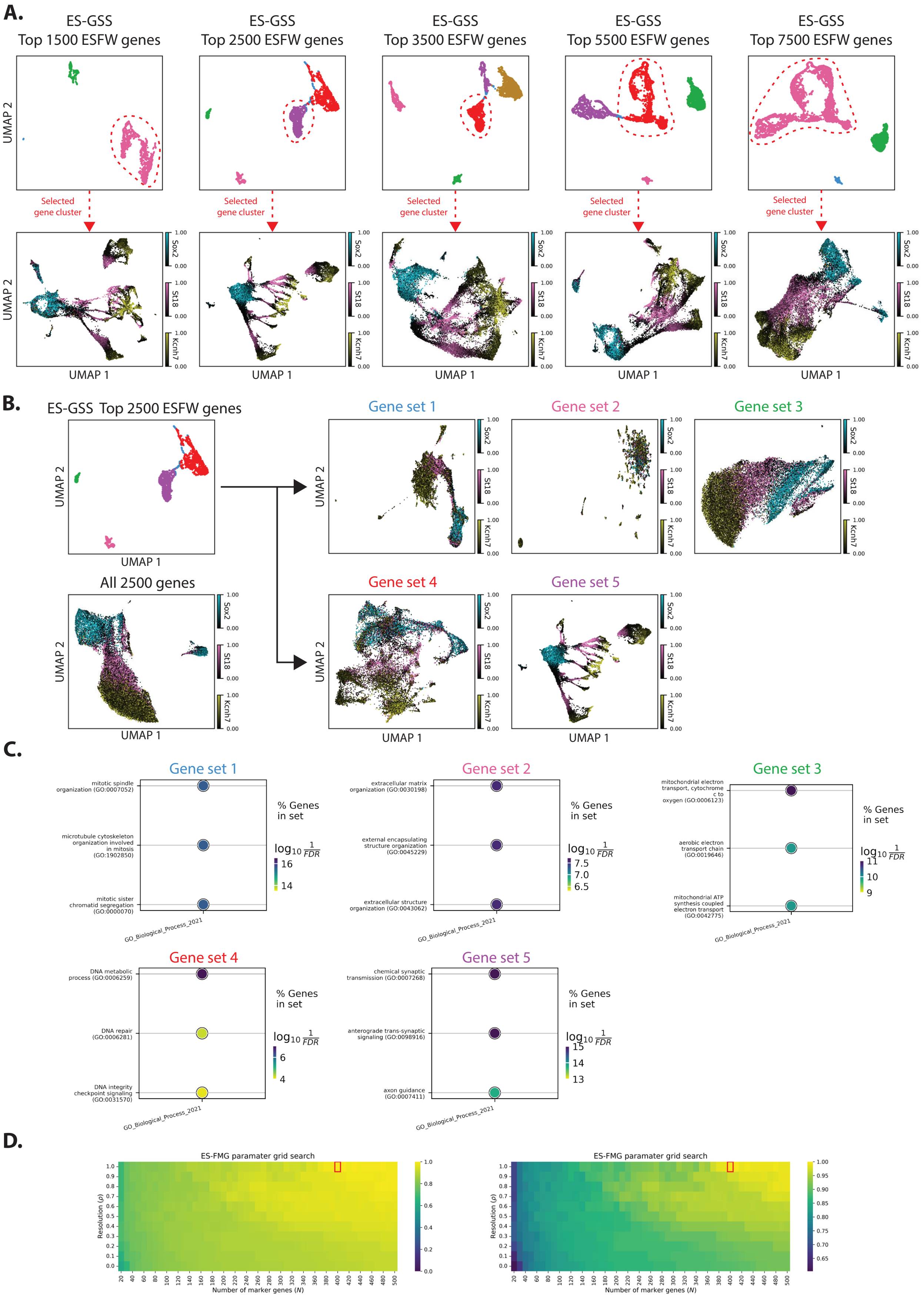


Figure S14: **ESFS parameter selection sensitivity analysis - Mouse neural tube scRNA-seq. A.** Visualisation of the effect of changing the number of top ranked genes used during gene clustering on the resulting cell UMAP embedding. Gene rankings are determined via entropy sorting feature weighting. **B.** Visualisation of gene cluster choice on the resulting cell UMAP embedding for the number of top ranked genes used in the main manuscript. **C.** Top gene ontology terms obtained via gene set enrichment analysis of biological processes for each of the gene clusters presented in B.. **D.** Sensitivity analysis of parameter selection for the ES-FMG algorithm. A grid search of the number of marker genes to optimise for $(N)$ and the resolution parameter $(\rho)$ shows the expression profiles captured quickly plateau to values close to 1. Red box indicates parameters used in the main manuscript.
